# Dynamically evolving novel overlapping gene as a factor in the SARS-CoV-2 pandemic

**DOI:** 10.1101/2020.05.21.109280

**Authors:** Chase W. Nelson, Zachary Ardern, Tony L. Goldberg, Chen Meng, Chen-Hao Kuo, Christina Ludwig, Sergios-Orestis Kolokotronis, Xinzhu Wei

## Abstract

Understanding the emergence of novel viruses requires an accurate and comprehensive annotation of their genomes. Overlapping genes (OLGs) are common in viruses and have been associated with pandemics, but are still widely overlooked. We identify and characterize *ORF3d*, a novel OLG in SARS-CoV-2 that is also present in Guangxi pangolin-CoVs but not other closely related pangolin-CoVs or bat-CoVs. We then document evidence of *ORF3d* translation, characterize its protein sequence, and conduct an evolutionary analysis at three levels: between taxa (21 members of *Severe acute respiratory syndrome-related coronavirus*), between human hosts (3978 SARS-CoV-2 consensus sequences), and within human hosts (401 deeply sequenced SARS-CoV-2 samples). *ORF3d* has been independently identified and shown to elicit a strong antibody response in COVID-19 patients. However, it has been misclassified as the unrelated gene *ORF3b*, leading to confusion. Our results liken *ORF3d* to other accessory genes in emerging viruses and highlight the importance of OLGs.

## Introduction

The COVID-19 pandemic raises urgent questions about the properties that allow animal viruses to cross species boundaries and spread within humans. Addressing these questions requires an accurate and comprehensive understanding of viral genomes. One frequently overlooked source of novelty is the evolution of new overlapping genes (OLGs), wherein a single stretch of nucleotides encodes two distinct proteins in different reading frames. Such “genes within genes” compress genomic information and allow genetic innovation via *overprinting* (Keese and Gibbs 1992), particularly as frameshifted sequences preserve certain physicochemical properties of proteins (Bartonek et al. 2020). However, OLGs also entail the cost that a single mutation may alter two proteins, constraining evolution of the pre-existing open reading frame (ORF) and complicating sequence analysis. Unfortunately, genome annotation methods typically miss OLGs, instead favoring one ORF per genomic region (Warren et al. 2010).

OLGs are known entities but remain inconsistently reported in viruses of the species *Severe acute respiratory syndrome-related coronavirus* (subgenus *Sarbecovirus*; genus *Betacoronavirus*; Gorbalenya et al. 2020). For example, annotations of *ORF3b*, *ORF9b*, and *ORF9c* are absent or conflicting in SARS-CoV-2 reference genome Wuhan-Hu-1 (NCBI: NC_045512.2) and genomic studies (e.g., Chan et al. 2020; F. Wu et al. 2020), and OLGs are often not displayed in genome browsers (e.g., Flynn et al. 2020). Further, *ORF3b* within *ORF3a*, an extensively characterized OLG present in other species members including SARS-CoV (SARS-CoV-1) (McBride and Fielding 2012), has sometimes been annotated in SARS-CoV-2 even though it contains four early STOP codons in this virus. Such inconsistencies complicate research, because OLGs may play key roles in the emergence of new viruses. For example, in human immunodeficiency virus-1 (HIV-1), the novel OLG *asp* within *env* is actively expressed in human cells (Affram et al. 2019) and is associated with the pandemic M group lineage (Cassan et al. 2016).

### Novel overlapping gene candidates

To identify novel OLGs within the SARS-CoV-2 genome, we first generated a list of candidate overlapping ORFs in the Wuhan-Hu-1 reference genome (NCBI: NC_045512.2). Specifically, we used the codon permutation method of Schlub et al. (2018) to detect unexpectedly long ORFs while controlling for codon usage. One unannotated gene candidate, here named *ORF3d*, scored highly (p=0.0104), exceeding the significance of two known OLGs annotated in Uniprot (*ORF9b* and *ORF9c*/*ORF14* within *N*; https://viralzone.expasy.org/8996) (Figure 1, Figure 1—figure supplement 1, and Supplementary Table 1).

**Figure 1.**
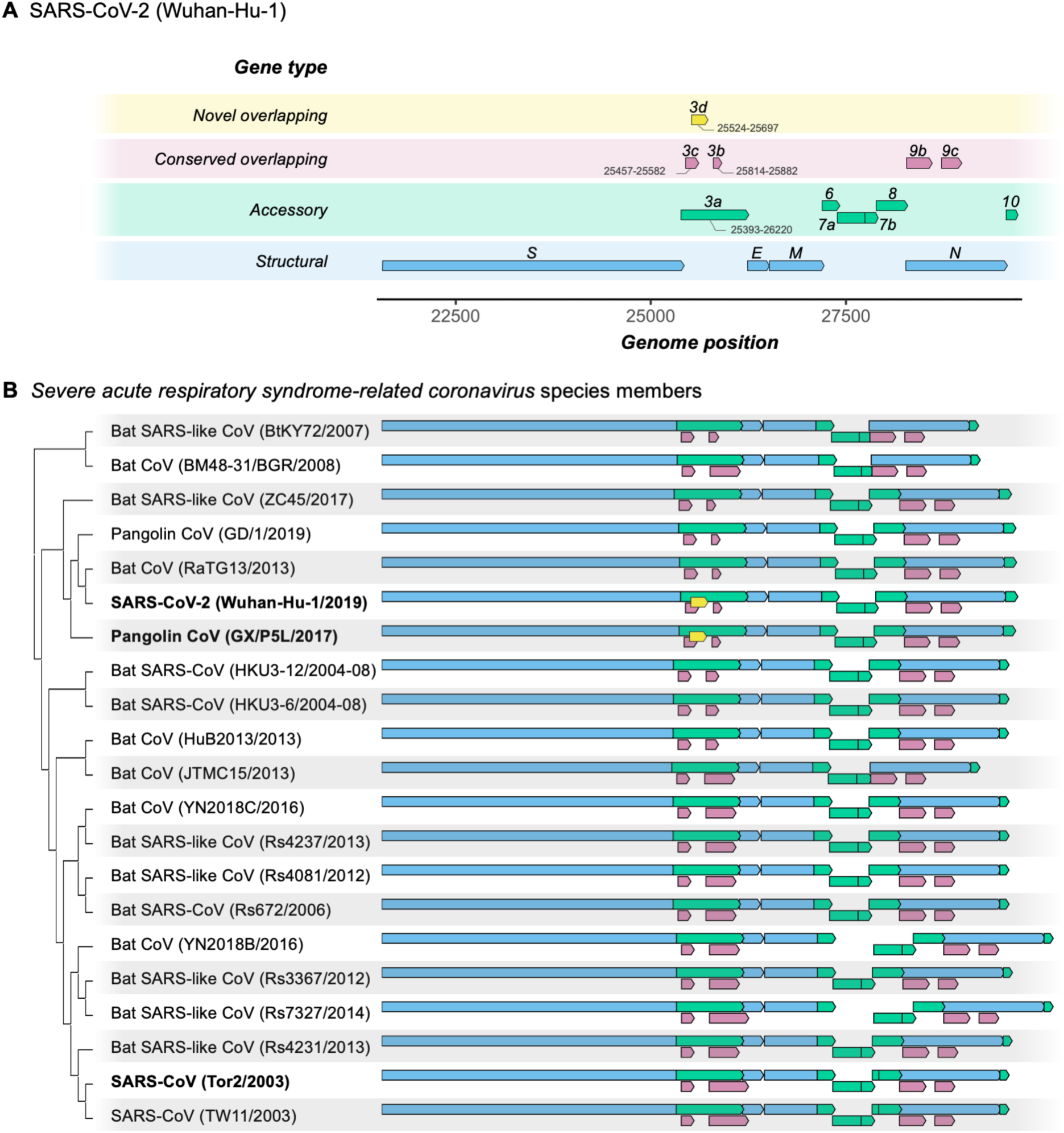
Gene repertoire and evolutionary relationships of *Severe acute respiratory syndrome-related coronavirus* species members. Only genes downstream of *ORF1ab* are shown, beginning with *S* (Spike-encoding). **(A)** Four types of genes and their relative positions in the SARS-CoV-2 Wuhan-Hu-1 genome (NCBI: NC_045512.2). Genes are colored by type: novel overlapping genes (OLGs) (gold; *ORF3d* only); conserved OLGs (burgundy); accessory (green); and structural (blue). Note that *ORF3b* has been truncated relative to SARS-CoV genomes, whereas *ORF8* remains intact (i.e., has not been split into *ORF8a* and *ORF8b*). **(B)** Genes with intact ORFs in each of the the 21 *Severe acute respiratory syndrome-related coronavirus* genomes. Gene positions are shown relative to each genome, i.e., homologous genes are not precisely aligned. Only the full-length versions of *ORF3a* and *ORF3d* are shown (for shorter isoforms, see Table 1). Note that the first 20 codons of *ORF3d* overlap the last codons of *ORF3c* (supplementary file SARS-CoV-2.gtf), such that the beginning of *ORF3d* involves a triple overlap (*ORF3a*/*ORF3c*/*ORF3d*). *ORF3b* is full-length in only 3 sequences (SARS-CoV TW11, SARS-CoV Tor2, and bat-CoV Rs7327), while the remaining sequences have either an early or a late premature STOP codon (Supplementary Table 9). *ORF8* is not novel in SARS-CoV-2 (contra Chan et al. 2020), but is intact in all but 5 sequences (split into *ORF8a* and *ORF8b* in SARS-CoVs TW11 and Tor2; deleted in bat-CoVs BtKY72, BM48-31, and JTMC15). *ORF9b* and *ORF9c* are found throughout this virus species, yet rarely annotated in genomes at NCBI. **Figure 1—source data 1.** SARS-related-CoV_ALN.fasta. Whole-genome multiple sequence alignment of 21 genomes of the species *Severe acute respiratory syndrome-related coronavirus*. Note that the pangolin-CoV GD/1 sequence has been masked as N (license constraint), but can be acquired with GISAID registration. **Figure 1—source data 2.** SARS-related-CoV_ALN.gtf.txt. Gene Transfer Format (GTF) file giving gene positions within SARS-related-CoV_ALN.fasta. Genes were extracted and gaps removed from each sequence before displaying.

**Table 1.**
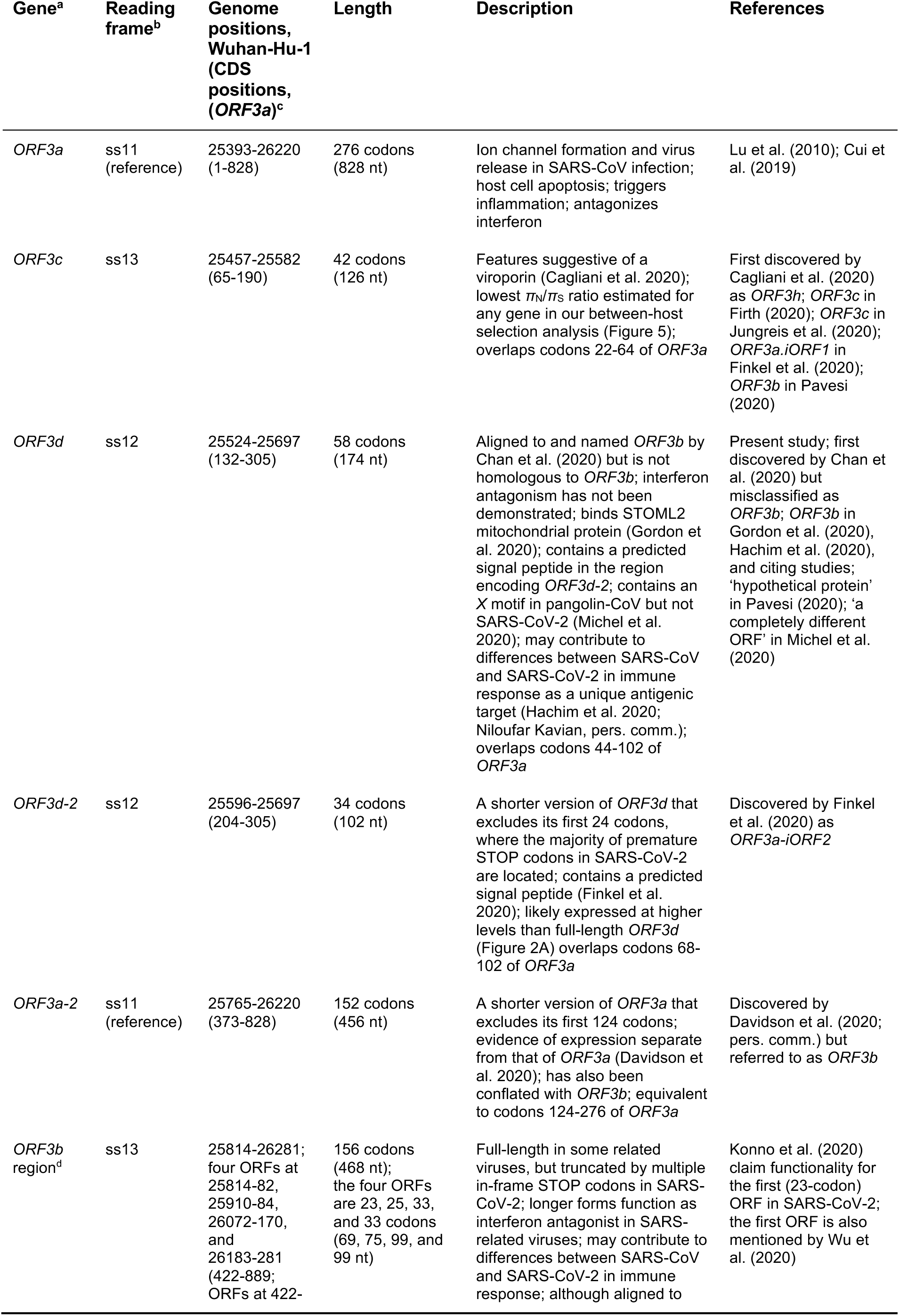
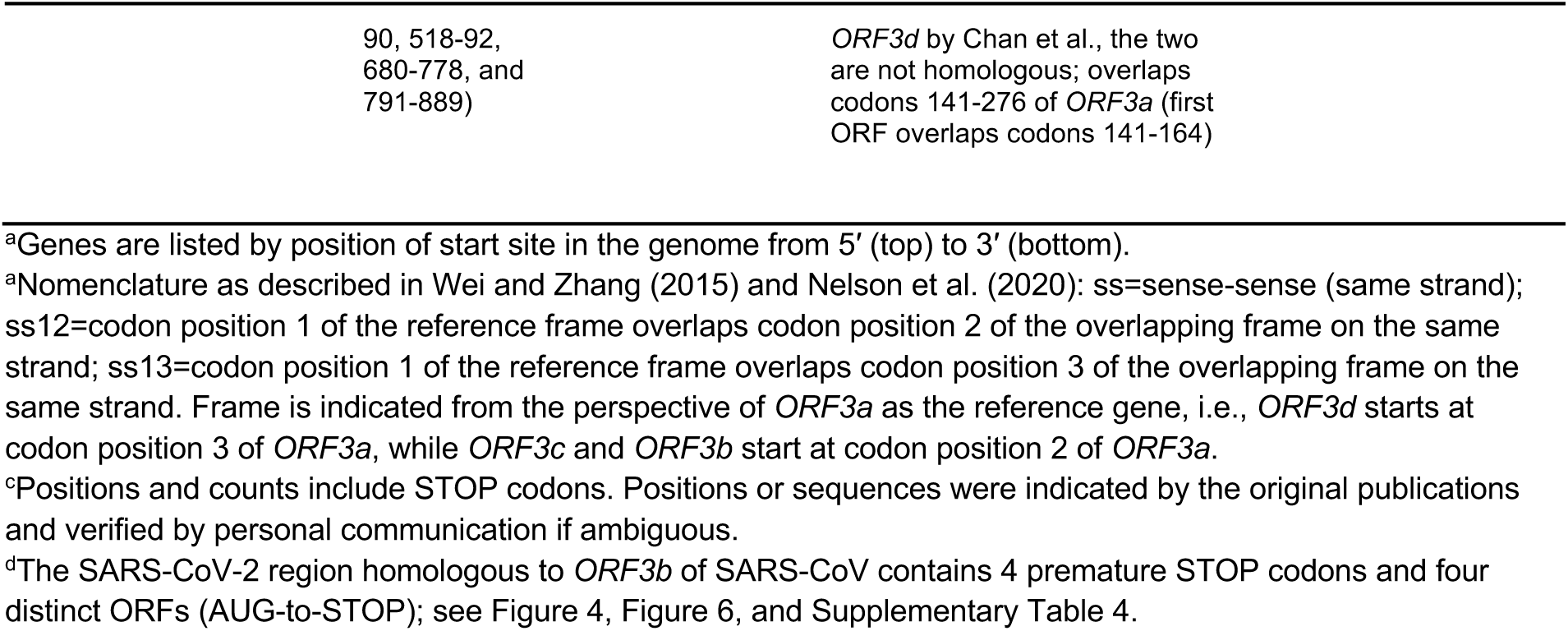
Nomenclature and reading frames for overlapping gene candidates in SARS-CoV-2 *ORF3a*.

| Gene<sup>a</sup> | Reading frame<sup>b</sup> | Genome positions, Wuhan-Hu-1 (CDS positions, *ORF3a*)<sup>c</sup> | Length | Description | References |
| --- | --- | --- | --- | --- | --- |
| *ORF3a* | ss11 (reference) | 25393-26220 (1-828) | 276 codons (828 nt) | Ion channel formation and virus release in SARS-CoV infection; host cell apoptosis; triggers inflammation; antagonizes interferon | Lu et al. (2010); Cui et al. (2019) |
| *ORF3c* | ss13 | 25457-25582 (65-190) | 42 codons (126 nt) | Features suggestive of a viroporin (Cagliani et al. 2020); lowest $\pi_N/\pi_S$ ratio estimated for any gene in our between-host selection analysis (Figure 5); overlaps codons 22-64 of *ORF3a* | First discovered by Cagliani et al. (2020) as *ORF3h*; *ORF3c* in Firth (2020); *ORF3c* in Jungreis et al. (2020); *ORF3a.iORF1* in Finkel et al. (2020); *ORF3b* in Pavesi (2020) |
| *ORF3d* | ss12 | 25524-25697 (132-305) | 58 codons (174 nt) | Aligned to and named *ORF3b* by Chan et al. (2020) but is not homologous to *ORF3b*; interferon antagonism has not been demonstrated; binds STOML2 mitochondrial protein (Gordon et al. 2020); contains a predicted signal peptide in the region encoding *ORF3d-2*; contains an X motif in pangolin-CoV but not SARS-CoV-2 (Michel et al. 2020); may contribute to differences between SARS-CoV and SARS-CoV-2 in immune response as a unique antigenic target (Hachim et al. 2020; Niloufar Kavian, pers. comm.); overlaps codons 44-102 of *ORF3a* | Present study; first discovered by Chan et al. (2020) but misclassified as *ORF3b*; *ORF3b* in Gordon et al. (2020), Hachim et al. (2020), and citing studies; 'hypothetical protein' in Pavesi (2020); 'a completely different ORF' in Michel et al. (2020) |
| *ORF3d-2* | ss12 | 25596-25697 (204-305) | 34 codons (102 nt) | A shorter version of *ORF3d* that excludes its first 24 codons, where the majority of premature STOP codons in SARS-CoV-2 are located; contains a predicted signal peptide (Finkel et al. 2020); likely expressed at higher levels than full-length *ORF3d* (Figure 2A) overlaps codons 68-102 of *ORF3a* | Discovered by Finkel et al. (2020) as *ORF3a.iORF2* |
| *ORF3a-2* | ss11 (reference) | 25765-26220 (373-828) | 152 codons (456 nt) | A shorter version of *ORF3a* that excludes its first 124 codons; evidence of expression separate from that of *ORF3a* (Davidson et al. 2020); has also been conflated with *ORF3b*; equivalent to codons 124-276 of *ORF3a* | Discovered by Davidson et al. (2020; pers. comm.) but referred to as *ORF3b* |
| *ORF3b* region<sup>d</sup> | ss13 | 25814-26281; four ORFs at 25814-82, 25910-84, 26072-170, and 26183-281 (422-889; ORFs at 422- | 156 codons (468 nt); the four ORFs are 23, 25, 33, and 33 codons (69, 75, 99, and 99 nt) | Full-length in some related viruses, but truncated by multiple in-frame STOP codons in SARS-CoV-2; longer forms function as interferon antagonist in SARS-related viruses; may contribute to differences between SARS-CoV and SARS-CoV-2 in immune response; although aligned to | Konno et al. (2020) claim functionality for the first (23-codon) ORF in SARS-CoV-2; the first ORF is also mentioned by Wu et al. (2020) |

<sup>a</sup>Genes are listed by position of start site in the genome from 5' (top) to 3' (bottom).
<sup>a</sup>Nomenclature as described in Wei and Zhang (2015) and Nelson et al. (2020): ss=sense-sense (same strand); ss12=codon position 1 of the reference frame overlaps codon position 2 of the overlapping frame on the same strand; ss13=codon position 1 of the reference frame overlaps codon position 3 of the overlapping frame on the same strand. Frame is indicated from the perspective of *ORF3a* as the reference gene, i.e., *ORF3d* starts at codon position 3 of *ORF3a*, while *ORF3c* and *ORF3b* start at codon position 2 of *ORF3a*.
<sup>c</sup>Positions and counts include STOP codons. Positions or sequences were indicated by the original publications and verified by personal communication if ambiguous.
<sup>d</sup>The SARS-CoV-2 region homologous to *ORF3b* of SARS-CoV contains 4 premature STOP codons and four distinct ORFs (AUG-to-STOP); see Figure 4, Figure 6, and Supplementary Table 4.

*ORF3d* comprises 58 codons (including STOP) near the beginning of *ORF3a* (Table 1), making it longer than the known genes *ORF7b* (44 codons) and *ORF10* (39 codons) (Supplementary Tables 2 and 3). *ORF3d* was discovered independently by Chan et. al (2020) as ‘*ORF3b*’ and Pavesi (2020) as, simply, ‘hypothetical protein’. Due to this naming ambiguity and its location within *ORF3a*, *ORF3d* has subsequently been conflated with the previously documented *ORF3b* in multiple studies (Fung et al. 2020; Ge et al. 2020; Gordon et al. 2020; Hachim et al. 2020; Helmy et al. 2020; Yi et al. 2020). Critically, *ORF3d* is unrelated (i.e., not homologous) to *ORF3b*, as the two genes occupy different genomic positions within *ORF3a*: *ORF3d* ends 39 codons upstream of the genome region homologous to *ORF3b*, and the *ORF3b* start site corresponds to an ORF encoding only 23 codons in SARS-CoV-2 (A. Wu et al. 2020) (Table 1, Figure 1, Figure 1—figure supplement 1, and Supplementary Table 4). Furthermore, the two genes occupy different reading frames: codon position 1 of *ORF3a* overlaps codon position 2 of *ORF3d* (frame ss12) but codon position 3 of *ORF3b* (frame ss13). *ORF3d* is also distinct from other OLGs hypothesized within *ORF3a* (Table 1). Thus, *ORF3d* putatively encodes a novel protein not present in other *Severe acute respiratory syndrome-related coronavirus* genomes, while the absence of full-length *ORF3b* in SARS-CoV-2 distinguishes it from SARS-CoV (Figure 1). Because *ORF3b* plays a central role in SARS-CoV immune interactions, its absence or truncation in SARS-CoV-2 may be immunologically important (Konno et al. 2020; Yuen et al. 2020).

### *ORF3d* molecular biology and expression

To assess the expression of *ORF3d*, we re-analyzed the SARS-CoV-2 ribosome profiling (Ribo-seq) data of Finkel et al. (2020). This approach allows the study of gene expression and reading frame at single nucleotide (nt) resolution by sequencing mRNA fragments bound by actively translating ribosomes (ribosome footprints) (Ingolia et al. 2009). Focusing on samples with reads reliably associated with ribosomes stalled at translation start sites (i.e., lactimidomycin and harringtonine treatments at 5 hours post-infection) and trimming reads to their 5′ ends (first nt), we observe a consistent peak in read depth ∼12 nt upstream of the start site of *ORF3d* (Figure 2A), the expected ribosomal P-site offset (Calviello and Ohler 2017). Similar peaks are also observed for previously annotated genes (Figure 2A, Figure 2—figure supplement 1).

**Figure 2.**
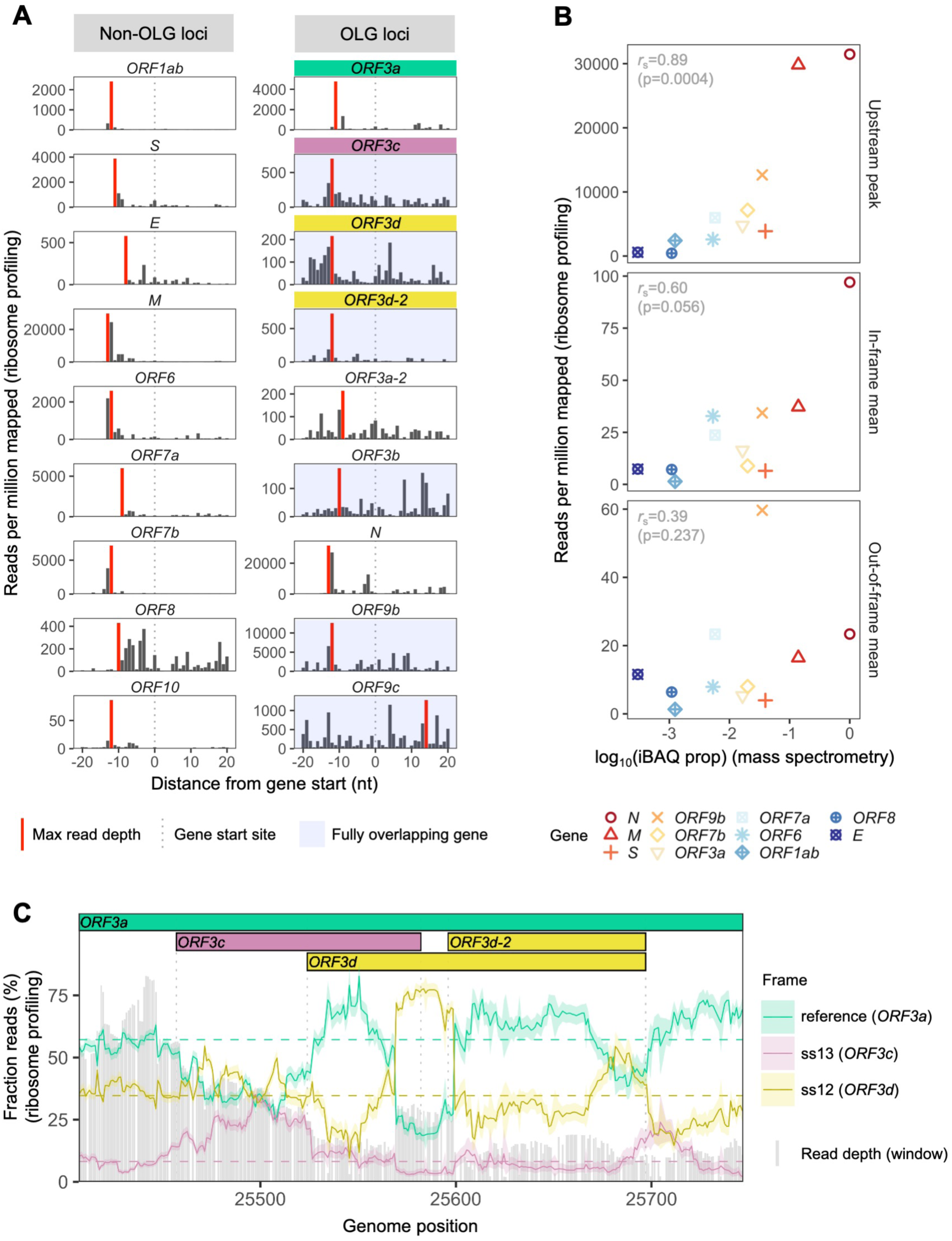
SARS-CoV-2 gene expression in ribosome profiling and mass spectrometry datasets. Ribosome profiling (Ribo-seq) data were obtained from Finkel et al. (2020); mass spectrometry data were obtained from Davidson et al. (2020) and Bezstarosti et al. (2020). Reads were trimmed to their first (5′) nucleotide to minimize statistical dependence while preserving reading frame. Results are shown after pooling samples treated with harringtonine (SRR11713360, SRR11713361) or lactimidomycin (SRR11713358, SRR11713359) at 5 hours post-infection. (**A**) Ribosome profiling coverage (read depth) near translation initiation sites, measured as mean reads per million mapped reads. Only reads of length 29-31 nucleotides were used, chosen for their enrichment at the start sites of highly expressed annotated genes (Figure 2—figure supplement 4). Light blue backgrounds denote fully (internal) overlapping genes. Annotated genes show an accumulation of 5′ ends of protected reads upstream of the gene’s start site (grey dotted vertical line), peaking near the ribosome P-site offset of -12 nt (red=maximum depth). Distributions are largely consistent across individual samples (Figure 2—figure supplement 1). The ranges of the y axes vary according to expression level, with the most highly expressed gene (*N*; Figure 2—figure supplement 2) having the largest range. (**B**) Correlation between protein expression as estimated by mass spectrometry and ribosomal profiling. ‘iBAQ prop’ refers to the relative (proportion of maximum) protein intensity-based absolute quantification (iBAQ) value (Schwanhäusser et al. 2011). Genes are denoted by shape and ordinally colored by iBAQ prop from high (red) to low (blue). ‘Upstream peak’ refers to the maximum read depth observed at the approximate P-site offset (red bars in Figure 2A), while mean read depths were measured across each gene using 30 nt reads separately for in-frame (codon position 1) and out-of-frame (codon positions 2 and 3) sites (non-overlapping gene regions only, except for *ORF9b*). (**C**) Reading frame of ribosome profiling reads in the *ORF3c*/*ORF3d* region of *ORF3a*. Solid lines show the fraction of reads in each frame, summed across samples in sliding windows of 30 nt (step size=1 nt; read length=30 nt). Color denotes frame: green=reference frame (*ORF3a*); burgundy=ss13, the sense frame encoding *ORF3c*, whose codon position 3 overlaps codon position 1 of *ORF3a*; and gold=ss12, the sense frame encoding *ORF3d*, whose codon position 2 overlaps codon position 1 of *ORF3a*. Values are shown for the central nucleotide of each window, with shaded regions corresponding to 95% binomial confidence intervals. Alternative frame translation is suggested where a given frame (solid line) exceeds its average across the remainder of the gene (horizontal dashed line; non-OLG regions of *ORF3a*). Vertical gray dotted lines indicate gene start and end sites. Gray bars show read depth for each window, with a maximum of 2889 reads at genome position 25442. **Figure 2—source data 1.** riboseq_upstream_peaks.txt. Source data for Figure 2A. **Figure 2—source data 2.** expression_data_by_gene_frame.txt. Source data for Figure 2B. **Figure 2—source data 3.** riboseq_ORF3d_sliding_window.txt. Source data for Figure 2C.

To investigate the relationship between ribosome profiling read depth and expressed protein levels, we next re-analyzed five publicly available SARS-CoV-2 mass spectrometry (MS) datasets (Bezstarosti et al. 2020; Bojkova et al. 2020; Davidson et al. 2020; PRIDE Project PXD018581; Zecha et al. 2020; Methods). We were unable to detect ORF3c, ORF3d, ORF3b, ORF9c, or ORF10 when employing a 1% false-discovery threshold. This result may reflect the limitations of MS for detecting proteins that are very short, weakly expressed under the specific conditions tested, or lack detectable peptides. Even the envelope protein E is not detected in some SARS-CoV-2 studies (Bojkova et al. 2020; Davidson et al. 2020), and the only evidence for ORF3b expression in SARS-CoV comes from Vero E6 cells (McBride and Fielding 2012). However, we do find a strong correlation between protein expression as estimated from MS and the ribosome profiling read depth observed at upstream peaks, with *r*_S_=0.89 (p=0.0004, Spearman’s rank) (Figure 2, Figure 2—figure supplement 2). This suggests that the presence and depth of an upstream peak is a reliable indicator of expression. Specifically, results from both proteomic and ribosome profiling approaches confirm that N, M, and S are the most highly expressed proteins, with N constituting ∼80% of the total viral protein content. We also observe that the number of reads determined to be in the correct reading frame (codon position 1) along the gene is moderately correlated with the MS expression values (*r*_S_=0.60, p=0.056), while this is not true for out-of-frame (codon position 2 and 3) reads (*r*_S_=0.39, p=0.237) (Figure 2B, Figure 2—figure supplement 2).

Ribosome profiling reads have a strong tendency to map with their first nucleotide occupying a particular codon position (i.e., in-frame) (Figure 2—figure supplement 3). We therefore examined codon position mapping across *ORF3a* to explore whether the proportions of reads in alternative frames are higher in the *ORF3c*/*ORF3d* region. Analyses were limited to reads of length 30 nt, the expected size of ribosome-bound fragments, which exhibit positioning indicative of the correct reading frame (codon position 1) of annotated genes and the highest total coverage (Figure 2—figure supplement 3, Figure 2—figure supplement 4). Sliding windows reveal a peak in the fraction of reads mapping to the reading frame of *ORF3d* at its hypothesized locus, with this frame’s maximum occurring at the center of *ORF3d*, co-located with a dip in the fraction of reads mapping to the frame of *ORF3a* (Figure 2C). These observations are reproducible across treatments (Figure 2— figure supplement 5), robust to sliding window size (Figure 2—figure supplement 6), and similar to the reading frame perturbations observed for the OLG *ORF9b* within *N* (Figure 2— figure supplement 7). At the same time, the *ORF3d* peak stands in contrast to the remainder of *ORF3a*, where the reading frame of *ORF3a* predominates, and smaller peaks in the frame of *ORF3d* are not accompanied by a large dip in the frame of *ORF3a* (Figure 2—figure supplement 8). These observations suggest that *ORF3d* is actively translated. Similar conclusions can be drawn for *ORF3c*, *ORF3d-2*, and *ORF9b* (Figure 2C, Figure 2—figure supplement 7, Figure 2—figure supplement 9).

Additional experiments have also provided evidence of ORF3d translation. Gordon et al. (2020) used overexpression experiments to demonstrate that ORF3d (referred to as ‘ORF3b’; Table 1) can be stably expressed and that it interacts with the mitochondrial protein STOML2. Most compellingly, ORF3d, ORF8, and N elicit the strongest antibody responses observed in COVID-19 patient sera, with ORF3d sufficient to accurately diagnose the majority of COVID-19 cases (Hachim et al. 2020). Because *ORF3d* is restricted to SARS-CoV-2 and pangolin-CoV (see below), this finding is unlikely to be due to cross-reactivity with another coronavirus, and provides strong independent evidence of *ORF3d* expression during infection.

### Protein sequence properties

To further investigate the antigenic properties of ORF3d, we predicted linear T-cell epitopes for each SARS-CoV-2 protein. We employed NetMHCpan (Jurtz et al. 2017) for MHC class I (cytotoxic CD8+ T-cells), an approach shown to accurately predict SARS-CoV-2 epitopes shared with SARS-CoV (Grifoni et al. 2020), and NetMHCIIpan (Reynission et al. 2020) for MHC class II (helper CD4+ T-cells). Specifically, we tested all 9 amino acid (MHC I) or 15 amino acid (MHC II) substrings of each viral protein for predicted weak or strong binding by MHC. Epitope density was estimated as the mean number of predicted epitopes per residue for each protein. We also tested two sets of negative controls: (1) randomized peptides generated from each protein, representing the result expected given amino acid content alone; and (2) short unannotated ORFs present in the SARS-CoV-2 genome, representing the result expected for real ORFs that have been evolving without functional constraint.

For CD8+ T cells, the lowest predicted epitope density occurs in ORF3d (1.5 per residue), which is unexpected given its own amino acid content (p=0.150) and compared to short unannotated ORFs (p=0.078; permutation tests). The next lowest densities occur in N and ORF8 (Figure 3A). Intriguingly, as previously mentioned, these three peptides (ORF3d, N, and ORF8) also elicit the strongest antibody (B-cell epitope) responses measured in COVID-19 patient sera (Hachim et al. 2020), suggesting a possible balance between CD8+ T and B cell epitopes. For CD4+ T cells, again, ORF3d has one of the lowest predicted epitope densities (5.6 per residue; p≥0.291), with lower values seen in only three other genes. However, focusing instead on the shorter ORF3d-2 isoform, this protein contains zero predicted epitopes, which is a highly significant depletion given its own amino acid content (p=0.001) and compared to short unannotated ORFs (p=0.001). These observations suggest either a predisposition towards immune escape, allowing gene survival, or the action of selective pressures on *ORF3d* or *ORF3d-2* to remove epitopes. In stark contrast to ORF3d, ORF3c has the highest observed density (8.4 per residue), apparently as a function of its highly constrained amino acid content (p=0.996) (see below). The enrichment of predicted epitopes in unannotated proteins (e.g., ORF3c and randomized peptides) but not N, for which numerous epitopes are annotated at the Immune Epitope Database (Grifoni et al. 2020), demonstrates that ascertainment or methodological biases cannot account for the depletion of predicted epitopes in ORF3d.

**Figure 3.**
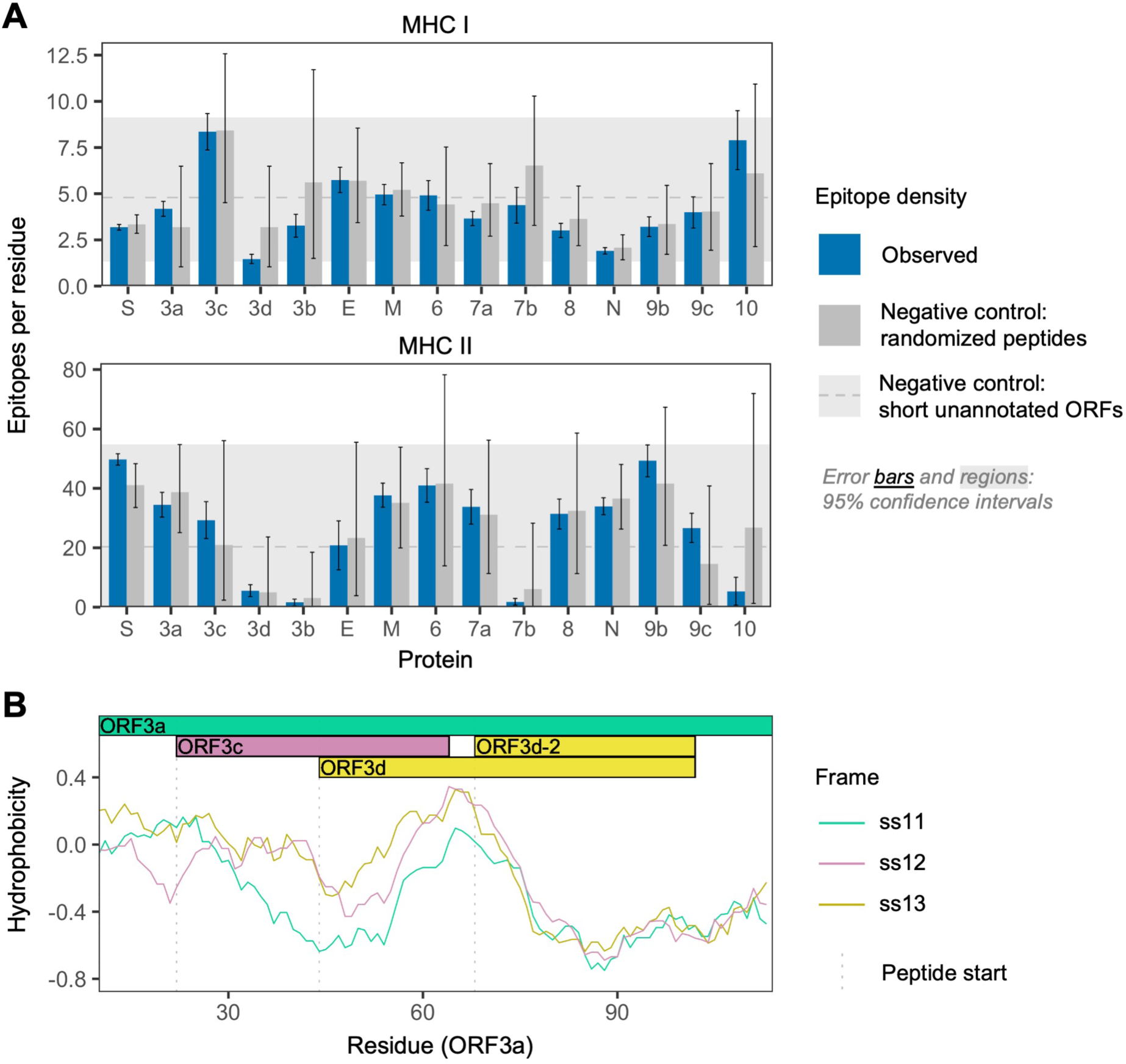
SARS-CoV-2 protein sequence properties. **(A)** Predicted densities of MHC class I-bound CD8+ T cell 9 aa epitopes (top) and MHC class II-bound CD4+ T cell 15 aa epitopes (bottom). Results for proteins encoded downstream of *ORF1ab* are shown. Mean numbers of predicted epitopes per residue (blue bars) are calculated as the number of epitopes overlapping each amino acid position divided by protein length. Error bars show 95% confidence intervals. Two sets of negative controls were also tested: (1) *n*=1,000 randomized peptides generated from each protein by randomly sampling its amino acids with replacement (dark gray bars), representing the result expected given amino acid content alone; and (2) short unannotated ORFs, the peptides encoded by *n*=103 putatively nonfunctional ORFs present in the SARS-CoV-2 genome, representing the result expected for real ORFs that have been evolving without functional constraint. For the short unannotated ORFs, the horizontal gray dotted line shows the mean number of epitopes per residue, and the gray shaded region shows the 95% confidence interval (*i.e.*, 2.5% to 97.5% quantiles). ORF3d and N have the lowest MHC class I epitope density; ORF3d, ORF3b, ORF7b, and ORF10 have the lowest MHC class II epitope density. **(B)** Hydrophobicity profiles of amino acid sequences in the three forward-strand reading frames of the *ORF3a*/*ORF3c*/*ORF3d* gene region, as calculated by the VOLPES server, using the unitless “Factor 1” consensus hydrophobicity scale. Frame is reported using *ORF3a* as the reference, e.g., ss12 refers to the frame encoding *ORF3d*, for which codon position 2 overlaps codon position 1 of *ORF3a* (Figure 2C). **Figure 3—source data 1.** epitope_summary_MHCI_LONG.txt. Source data for MHC class I in Figure 3A. **Figure 3—source data 2.** epitope_summary_MHCII_LONG.txt. Source data for MHC class II in Figure 3A. **Figure 3—source data 3.** hydrophobicity_profiles_ORF3a_corr.txt. Source data for hydrophobicity profiles in Figure 3B.

Our structural prediction for the ORF3d protein suggests *α*-helices connected with coils and an overall fold model that matches known protein structures (e.g., Protein Data Bank IDs 2WB7 and 6A93) with borderline confidence (Figure 3—figure supplement 1), similar to the predictions of Chan et al. (2020). Remarkably, biochemical properties that influence the structure of this novel protein appear to be inherited from the pre-existing ORF3a protein sequence encoded by the overlapping alternate reading frame. It has recently been shown that frame-shifted nucleotide sequences tend to encode proteins with similar hydrophobicity profiles as a consequence of the standard genetic code (Bartonek et al., 2020). To explore whether this is the case for *ORF3d*, we used the VOLPES server (Bartonek and Zagrovic 2019) to calculate the hydrophobicity profiles (Atchley et al. 2005) of the peptides encoded by all three frames of *ORF3a*. The maximum correlation observed occurs between the frames of ORF3a (ss11) and ORF3d (ss12) in the region encoding ORF3d (ORF3a residues 64-102), with *r*_S_=0.87 (p=5.79×10^-13^, Spearman’s rank), much stronger than the correlation between these frames observed for the non-OLG residues of ORF3a (*r*_S_=0.27, p=8.70×10^-4^) (Figure 3C; Figure 3—figure supplement 2). This conservation of a structure-related property with the ORF3a protein provides further evidence for *ORF3d* functionality, and may predispose this region towards *de novo* gene birth. Again, ORF3c shows the opposite pattern, as the minimum correlation observed between hydrophobicity profiles occurs between the frames of *ORF3a* (ss11) and *ORF3c* (ss13) in the region encoding ORF3c (ORF3a residues 22-43), with *r*_S_=-0.40 (p=6.16×10^-2^). Such disparate hydrophobicity profiles may be due to the strong conservation of *ORF3c* (see below).

### *ORF3d* taxonomic distribution and origins

To assess the origin of *ORF3d* and its conservation within and among host taxa, we aligned 21 *Severe acute respiratory syndrome-related coronavirus* genomes reported by Lam et al. (2020) (Supplementary Table 5), limiting our analysis to those with an annotated *ORF1ab* and no frameshift variants relative to SARS-CoV-2 in the core genes *ORF1ab*, *S*, *ORF3a*, *E*, *M*, *ORF7a*, *ORF7b*, and *N* (Supplementary Table 6; SARS-related-CoV_ALN.fasta, supplementary data). All other genes are also intact (i.e., there is no mid-sequence STOP codon) in all genomes (Supplementary Tables 7 and 8), with the exception of *ORF3d*, *ORF3b*, and *ORF8* (Figure 4). Specifically, *ORF3d* is intact in only 2 sequences: SARS-CoV-2 Wuhan-Hu-1 and pangolin-CoVs from Guangxi (GX/P5L). Full-length *ORF3b* is intact in only 3 sequences: SARS-CoV TW11, SARS-CoV Tor2, and bat-CoV Rs7327, with the remainder falling into two distinct groups sharing an early or late STOP codon, respectively (Supplementary Table 9). Finally, *ORF8* is intact in all but 5 sequences, where it contains premature STOP codons or large-scale deletions (Figure 1B).

**Figure 4.**
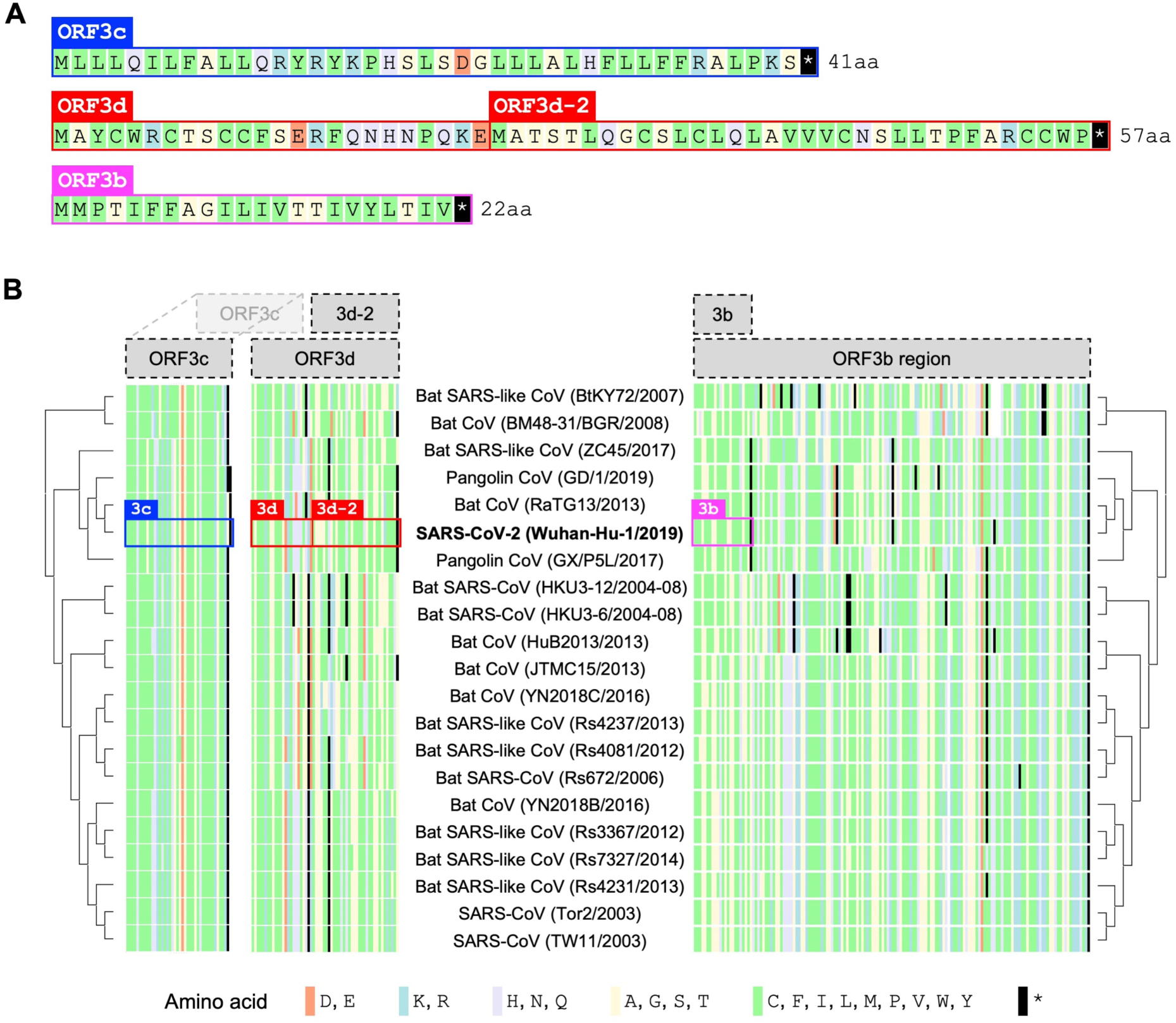
Amino acid variation in proteins encoded by genes overlapping *ORF3a* in viruses of the species *Severe acute respiratory syndrome-related coronavirus*. **(A)** Amino acid sequences of ORF3c, ORF3d, and ORF3b, as encoded by SARS-CoV-2 (Wuhan-Hu-1; NCBI: NC_045512.2). Note that, while ORF3b encodes a protein of 154 amino acids (aa) in SARS-CoV (NCBI: NC_004718.3), a premature STOP codon in SARS-CoV-2 has resulted in an ORF encoding only 22 aa. **(B)** Amino acid alignments of ORF3c, ORF3d, and ORF3b (3b) show their sequence conservation. Black lines indicate STOP codons in *ORF3d* and *ORF3b*, showing their restricted taxonomic ranges. Intact *ORF3d* is restricted to SARS-CoV-2 and pangolin-CoV GX/P5L; however, note that ORF3d-2 (denoted 3d-2), a shorter isoform of ORF3d, could have a slightly wider taxonomic range if TTG or GTG are permitted as translation initiation codons. Full-length ORF3b (ORF3b region) is found throughout members of this virus species, but truncated early in most genomes outside of SARS-CoV (Supplementary Tables 2, 3, 4, and 8), with the shortest version (denoted 3b) found in SARS-CoV-2 and closely related viruses. **Figure 4—source data 1.** aa_alignments_3c_3d_3b.xlsx. Amino acid alignments shown in Figure 4. The full translation of the *ORF3b* region is provided for ORF3b. Note that the pangolin-CoV GD/1 sequence has been masked as X (license constraint), but can be acquired with GISAID registration.

The presence of intact *ORF3d* homologs among viruses infecting different host species (human and pangolin) raises the possibility of functional conservation. However, the taxonomic distribution of this ORF is incongruent with whole-genome phylogenies in that *ORF3d* is intact in the pangolin-CoV more distantly related to SARS-CoV-2 (GX/P5L) but not the more closely related one (GD/1) (Figure 4), a finding confirmed by the alignment of Boni et al. (2020). New sequence data reveal similarly puzzling trends: *ORF3d* contains STOP codons in the closely related bat-CoV RmYN02 (GISAID: EPI_ISL_412977; data not shown), but it is intact in three more distantly related bat-CoVs discovered in Rwanda and Uganda, where it is further extended by 20 codons (total 78 codons) but shows no evidence of conservation with SARS-CoV-2 (Wells et al. 2020 and pers. comm; data not shown). Further, phylogenies of the 21 *Severe acute respiratory syndrome-related coronavirus* genomes built on *ORF3a* are incongruent with whole-genome phylogenies, likely due to the presence of recombination breakpoints in *ORF3a* near *ORF3d* (Boni et al. 2020; Rehman et al. 2020). Recombination, convergence, and recurrent loss may therefore each have played a role in the origin and taxonomic distribution of *ORF3d*.

### Between-taxa divergence

To examine natural selection on *ORF3d*, we measured viral diversity at three hierarchical evolutionary levels: between-taxa, between-host, and within-host. Specifically, between-taxa refers to divergence (*d*) among the 21 aforementioned viruses infecting bat, human, or pangolin (Figure 1B); between-host refers to diversity (*π*) between consensus-level SARS-CoV-2 genomes infecting different human individuals; and within-host refers to *π* in deeply sequenced ‘intrahost’ SARS-CoV-2 samples from single human individuals. At each level, we inferred selection by estimating mean pairwise nonsynonymous (*d*_N_ or *π*_N_; amino acid changing) and synonymous (*d*_S_ or *π*_S_; not amino acid changing) distances among all sequences. Importantly, we combined standard (non-OLG) methods (Nei and Gojobori 1986; Nelson et al. 2015) with OLGenie, a new *d*_N_/*d*_S_ method tailored for OLGs (hereafter OLG *d*_N_/*d*_S_), which we previously used to verify purifying selection on a novel OLG in HIV-1 (Nelson et al. 2020).

The only gene to show significant evidence of purifying selection at all three evolutionary levels is *N* (nucleocapsid-encoding gene) (Figure 5), which undergoes disproportionately low rates of nonsynonymous change (*d*_N_/*d*_S_<1 and *π*_N_/*π*_S_<1) specifically in its non-OLG regions, evidencing strict functional constraint. *N* is also the most highly expressed gene (Figure 2—figure supplement 2), confirming that selection has more opportunity to act when a protein is manufactured in abundance (Figure 2B; Supplementary Table 10) (Methods). Importantly, this signal can be missed if non-OLG methods are applied to *N* without accounting for its internal OLGs, *ORF9b* and *ORF9c* (e.g., at the between-host level, p=0.0268 when excluding OLG regions, but p=0.411 when including them; Supplementary Table 11).

**Figure 5.**
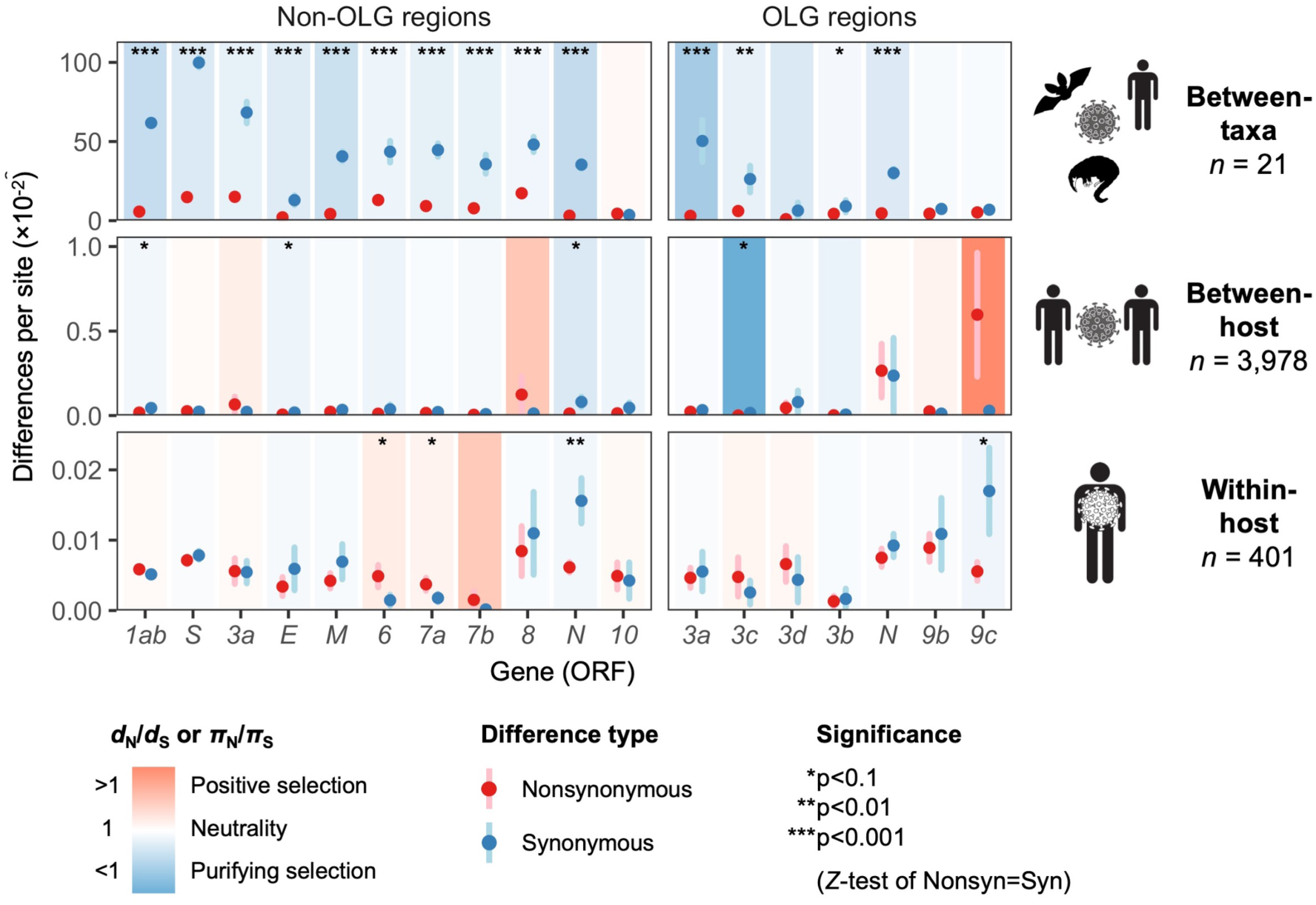
Natural selection analysis of viral nucleotide differences at three hierarchical evolutionary levels. Nucleotide differences in each virus gene were analyzed at three host levels: between-taxa divergence (*d*) among *Severe acute respiratory syndrome-related coronavirus* genomes infecting bat, human, and pangolin species; between-host diversity (*π*) for SARS-CoV-2 infecting human individuals (consensus-level); and within-host diversity (*π*) for SARS-CoV-2 infecting human individuals (deep sequencing). Each gene/level is shaded according to the ratio of mean nonsynonymous to synonymous differences per site to indicate purifying selection (*d*_N_/*d*_S_<1 or *π*_N_/*π*_S_<1; blue) or positive selection (*d*_N_/*d*_S_>1 or *π*_N_/*π*_S_>1; red), except the value for *ORF3c* was artificially lowered to allow the display of other ratios. A Jukes-Cantor correction was applied to *d*_N_/*d*_S_ values. Values range from a minimum of *π*_N_/*π*_S_=0.04 (*ORF3c*, between-host; p=0.0410, Q=0.308) to a maximum of 21.0 (*ORF9b*, between-host; p=0.126, Q=0.453), where significance (Q) was evaluated using a Benjamini-Hochberg false-discovery rate correction after *Z*-tests of the hypothesis that *d*_N_-*d*_S_=0 or *π*_N_-*π*_S_=0 (10,000 bootstrap replicates, codon unit). The mean of all pairwise comparisons is shown for sequenced genomes only, i.e., no ancestral sequences were reconstructed or inferred. For each gene, sequences were only included in the between-species analysis if a complete, intact ORF (no STOPs) was present. Genes containing an overlapping gene (OLG) in a different frame were analyzed separately for non-OLG and OLG regions using SNPGenie and OLGenie, respectively. For *ORF3b*, only the region corresponding to the first ORF in SARS-CoV-2 (Table 1) was analyzed. The short overlap between *ORF1a* and *ORF1b* (*nsp11* and *nsp12*) was excluded from the analysis. Error bars represent the standard error of mean pairwise differences. See Methods for further details. **Figure 5—source data 1.** selection_three_levels.txt. Source data for Figure 5.

With respect to *ORF3d*, comparison of Wuhan-Hu-1 to pangolin-CoV GX/P5L (NCBI: MT040335.1) yields OLG *d*_N_/*d*_S_=0.14 (p=0.264) (Figure 5), whereas inclusion of a third allele found in pangolin-CoV GX/P4L (NCBI: MT040333.1) yields 0.43 (p=0.488) (Supplementary Table 12). Because this is suggestive of constraint, we performed sliding windows of OLG *d*_N_/*d*_S_ across the length of *ORF3a* to check whether the evidence for purifying selection is specific to the expected genome comparisons and genome positions. Indeed, pairwise comparisons of each sequence to SARS-CoV-2 reveal OLG *d*_N_/*d*_S_<1 that is specific to the reading frame, genome positions, and species in which *ORF3d* is intact (pangolin-CoV GX/P5L) (Figure 6, left). This signal is independent of whether STOP codons are present, so its consilience with the only intact ORF in this region and species is highly suggestive of purifying selection. We note that this conclusion does not contradict studies which fail to find evidence of *ORF3d* conservation when comparing taxa where *ORF3d* is absent (e.g., Jungreis et al. 2020), because *ORF3d* is a novel gene and would by definition lack such conservation. This contrastive signal is also similar to what is observed for the known OLGs *ORF3b* (in comparisons to SARS-CoV; Figure 6, right), and *ORF9b* and *ORF9c* (in both viruses; Figure 6—figure supplement 1).

**Figure 6.**
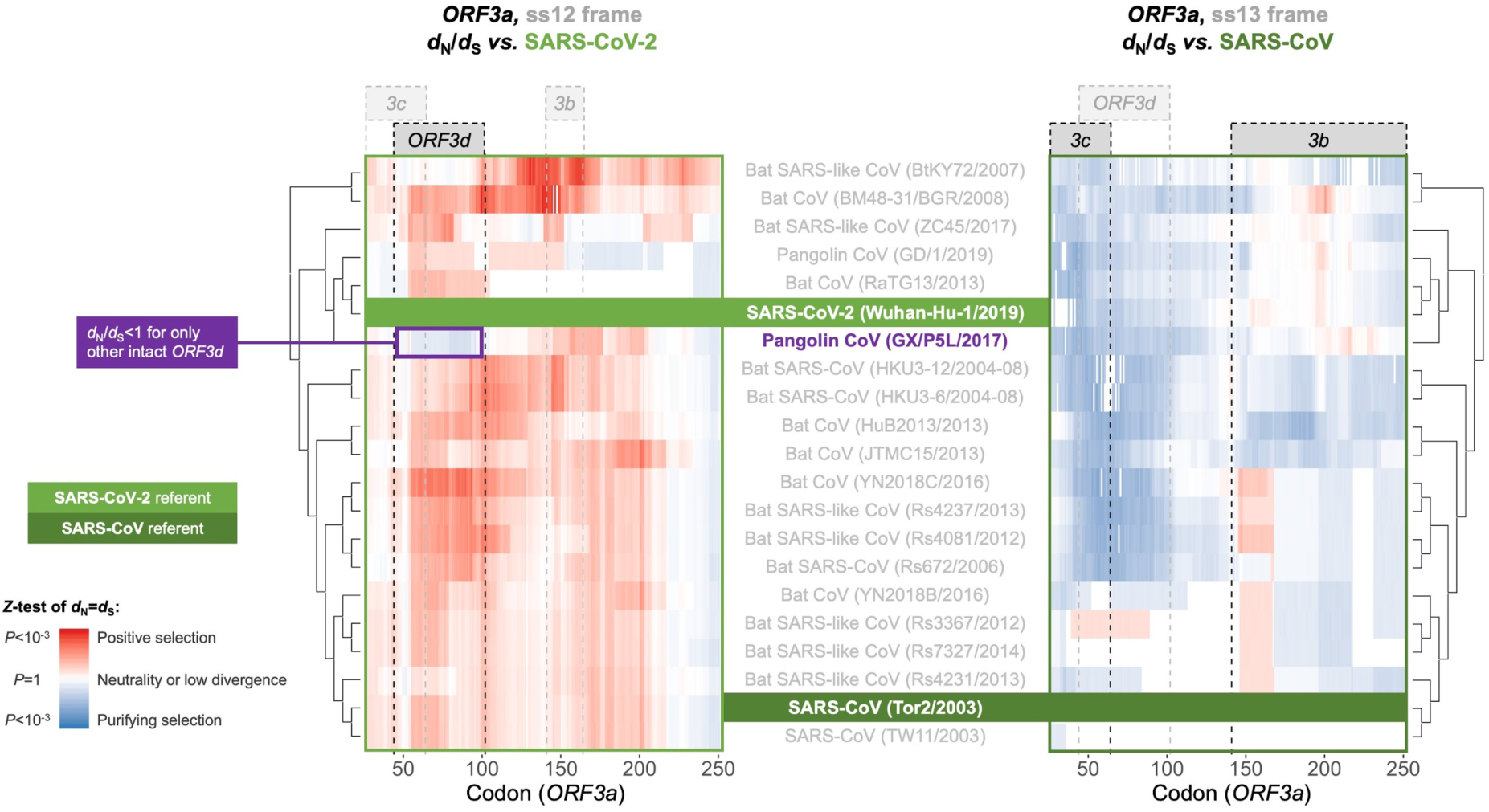
Between-taxa sliding window analysis of natural selection on overlapping frames of *ORF3a*. Pairwise sliding window analysis (windows size=50 codons; step size=1 codon) of selection across members of the species *Severe acute respiratory syndrome-related coronavirus*. OLG-appropriate *d*_N_/*d*_S_ values were computed using OLGenie (Nelson et al. 2020), a method that is conservative (non-conservative) for detecting purifying (positive) selection, and a Jukes-Cantor correction for multiple hits was employed. On the left-hand side, each genome is compared to SARS-CoV-2 in the ss12 reading frame of *ORF3a*, which contains *ORF3d* (Table 1 and Figure 2C). This frame shows evidence for purifying selection specific to the *ORF3d* region that is limited to the comparison with pangolin-CoV GX/P5L. On the right-hand side, this analysis is repeated for the ss13 reading frame of *ORF3a*, which contains *ORF3c* and *ORF3b* (Table 1 and Figure 2C), this time with respect to SARS-CoV, where full-length *ORF3b* is functional. This frame shows constraint across much of this gene and virus species, and that *ORF3c* in particular is deeply conserved. **Figure 6—source data 1.** SARS-CoV-2-ref_ORF3a_ss12_windows.txt. OLGenie sliding window *d*_N_/*d*_S_ results for comparisons to SARS-CoV-2 in the frame of *ORF3d*, shown on the left side of Figure 6. **Figure 6—source data 2.** SARS-CoV-ref_ORF3a_ss13_windows.txt. OLGenie sliding window *d*_N_/*d*_S_ results for comparisons to SARS-CoV in the frame of *ORF3b*, shown on the right side of Figure 6.

### Between-host evolution and pandemic spread

Purifying selection can be specific to just one taxon, as in the case of novel genes. Thus, to measure selection within SARS-CoV-2 only, we obtained *n*=3,978 high-quality human SARS-CoV-2 consensus sequences from GISAID (accessed April 10, 2020; Supplementary Table 13; GISAID Acknowledgments Table, supplementary data). Between-host diversity was sufficient to detect marginally significant purifying selection across all genes (*π*_N_/*π*_S_=0.50, p=0.0613, *Z*-test; Figure 5—figure supplement 2) but not most individual genes (Figure 5). Therefore, we instead investigated single mutations over time, limiting to 27 high-frequency variants (minor allele frequency ≥2%; Supplementary Table 14).

One high-frequency mutation occurred in *ORF3d*: G25563U, here denoted *ORF3d*-LOF (*ORF3d*-loss-of-function), which causes a STOP codon in *ORF3d* (ORF3d-E14*) but a nonsynonymous change in *ORF3a* (ORF3a-Q57H). This mutation is not observed in any other species member included in our analysis (Figure 4; SARS-related-CoV_ALN.fasta, supplementary data). During the first months of the COVID-19 pandemic, *ORF3d*-LOF increased in frequency (Supplementary Table 15) in multiple locations (Figure 5—figure supplement 2D; Supplementary Table 16), making this mutation a candidate for natural selection on ORF3a, ORF3d, or both (Kosakovsky-Pond 2020). However, temporal allele frequency trajectories (Figure 5—figure supplement 2D) and similar signals from phylogenetic branch tests may also be caused by founder effects or genetic drift, and are susceptible to ascertainment bias (e.g., preferential sequencing of imported infections and uneven geographic sampling) and stochastic error (e.g., small sample sizes).

To partially account for these confounding factors, we instead constructed the mutational path leading from the SARS-CoV-2 haplotype collected in December 2019 to the haplotype carrying *ORF3d*-LOF. This path involves five mutations (C241U, C3037U, C14408U, A23403G, G25563U), constituting five observed haplotypes (EP–3 → EP–2 → EP → EP+1 → EP+1+LOF), shown in Table 2. Here, EP is suggested to have driven the European Pandemic (detected in German patient #4; see footnote 3 of Table 2; Korber et al. 2020; Rothe et al. 2020); EP–3 is the Wuhan founder haplotype; EP–1 is never observed in our dataset; and LOF refers to *ORF3d*-LOF. We then documented the frequencies and earliest collection date of each haplotype (Table 2) to determine whether *ORF3d*-LOF occurred early on the EP background.

**Table 2.**
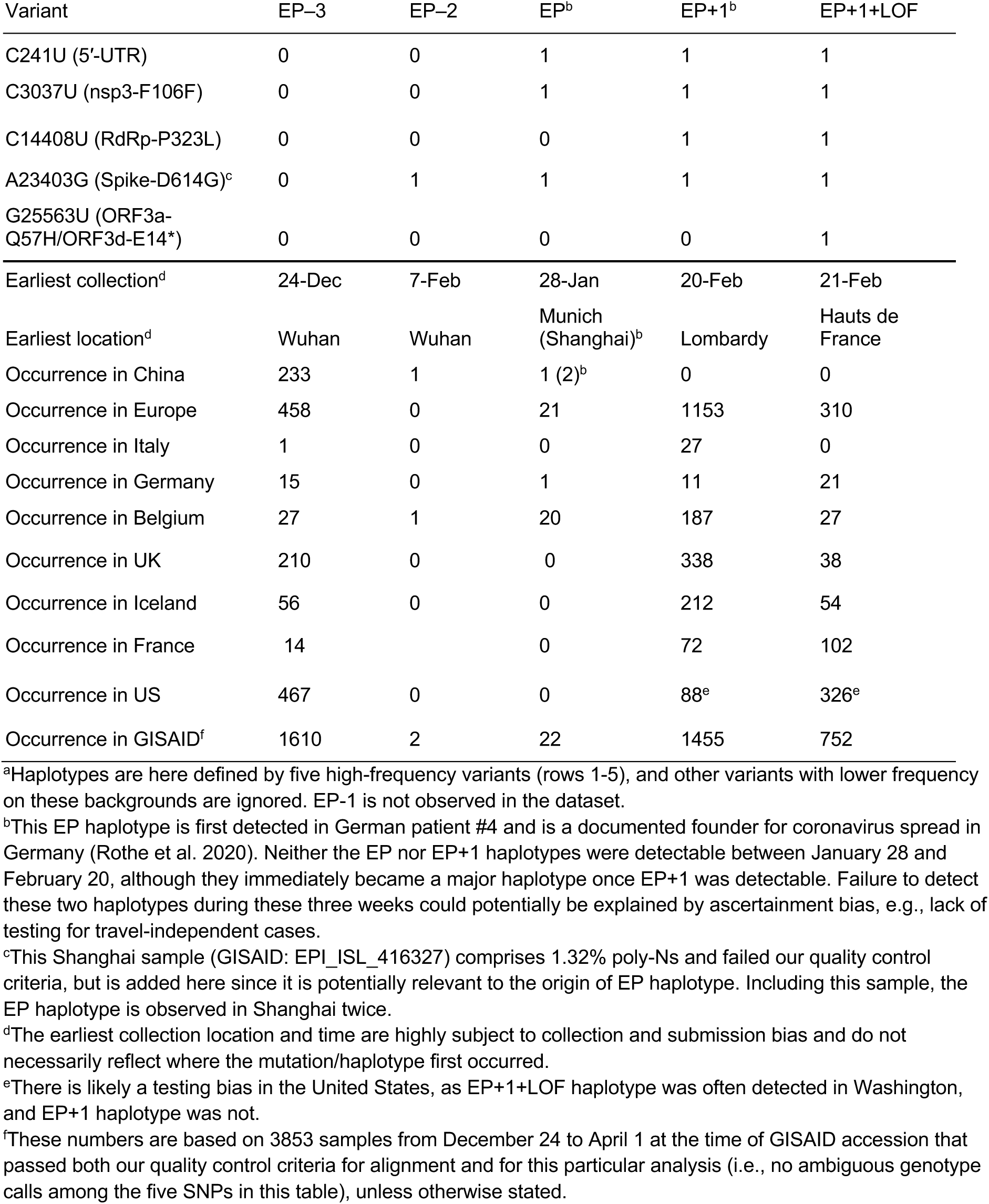
The mutational path to European pandemic founder haplotypes^a^

Surprisingly, despite its expected predominance in Europe due to a founder effect, the EP haplotype is extremely rare. By contrast, haplotypes with one additional mutation (C14408U; RdRp-P323L) on the EP background are common in Europe, and *ORF3d*-LOF occurred very early on this background to create EP+1+LOF from EP+1. Neither of these two haplotypes was initially observed in China (Table 2), suggesting that they might have arisen in Europe. Thus, we further partitioned the samples into two groups, corresponding to countries with early founders (January samples) or only late founders (no January samples) (Figure 7). In the early founder group, EP–3 (Wuhan) is the first haplotype detected in most countries, consistent with most early COVID-19 cases being related to travel from Wuhan. Because this implies that genotypes EP–3 and EP had longer to spread in the early founder group, it is surprising that their spread is dwarfed by the increase of EP+1 and EP+1+LOF starting in late February. This turnover is most evident in the late founder group, where multiple haplotypes are detected in a narrow time window, and the number of cumulative samples is always dominated by EP+1 and EP+1+LOF. Thus, while founder effects and drift are plausible explanations, it is also worth investigating whether the early spread of *ORF3d*-LOF may have been caused by its linkage with another driver, either C14408U (+1 variant) or a subsequent variant(s) occurring on the EP+1+LOF background (Discussion).

**Figure 7.**
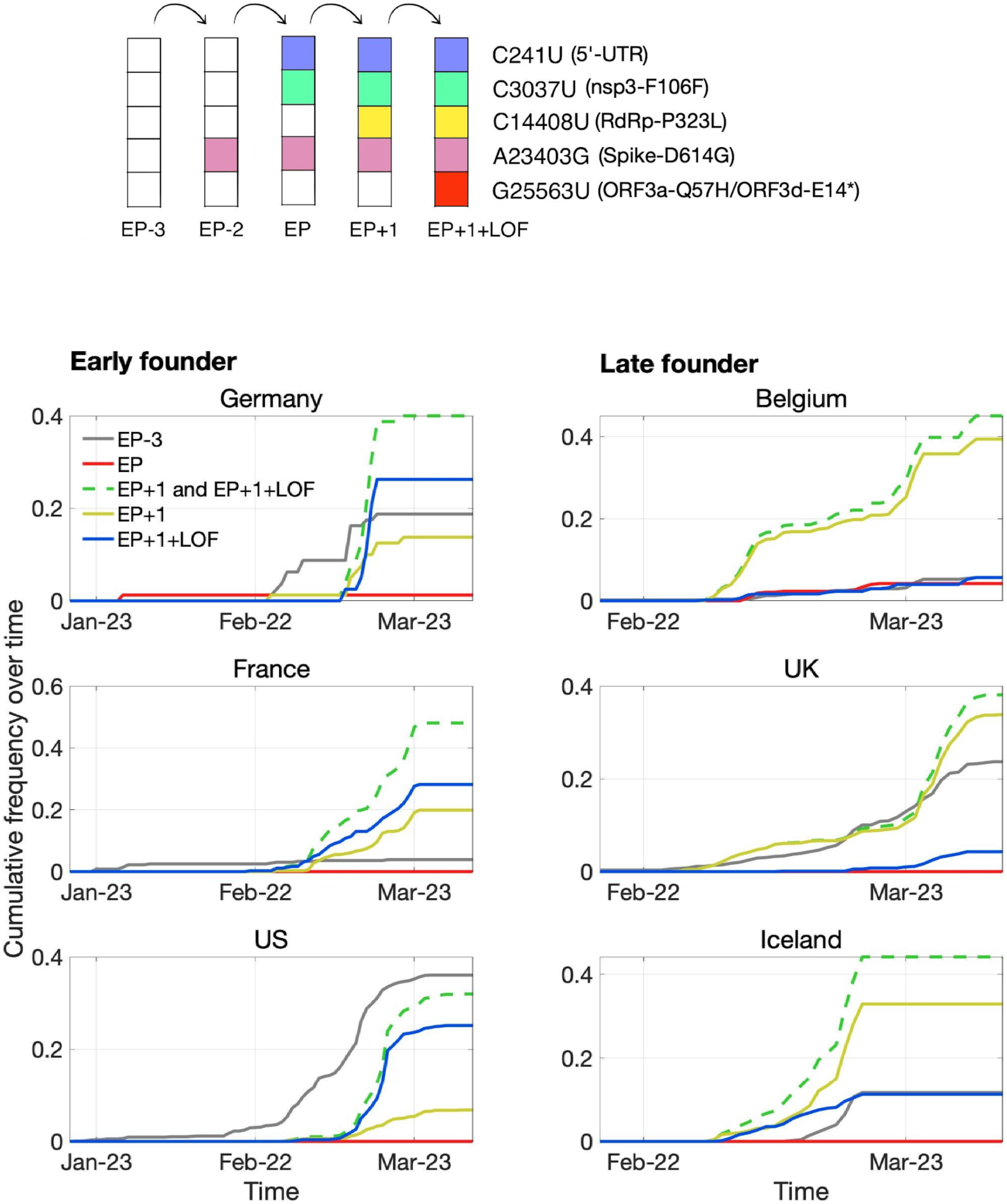
Pandemic spread of the EP+1 haplotype and the hitchhiking of *ORF3d*-LOF. The mutational path leading to EP+1+LOF is shown in the upper panel. Cumulative frequencies of haplotypes in samples from Germany and five other countries with the most abundant sequence data are shown in the lower panel. Countries are grouped into early founder (left) and late founder (right) based on the presence or absence of SARS-CoV-2 samples from January, respectively. In the early founder group, EP–3 (gray) is observed earlier than other haplotypes in France and the US, and EP (red) is observed early in Germany, giving them the advantage of a founder effect. However, neither EP nor EP–3 dominate later spread; instead, EP+1 (yellow) and EP+1+LOF (blue) increase much faster despite their later appearance in these countries. In the late founder group, multiple haplotypes appear at approximately the same time, but EP-3 and EP spread more slowly. The green dashed line shows the combined frequencies of EP+1 and EP+1+LOF (yellow and blue, respectively). Note that EP–1 is never observed in our dataset.

### Within-host diversity and mutational bias

Examination of within-host variation across multiple samples allows detection of recurrent mutations, which might indicate mutation bias or within-host selection. To investigate these possibilities, we obtained *n*=401 high-depth ‘intrahost’ human SARS-CoV-2 samples from the Sequence Read Archive (Supplementary Table 17) and called SNPs relative to the Wuhan-Hu-1 reference genome. Within human hosts, 42% of SNPs passed our false-discovery rate criterion (Methods), with a median passing minor allele frequency of 2% (21 of 1344 reads). Using these variants to estimate within-host diversity, *ORF3d* does not show significant evidence of selection, with OLG *π*_N_/*π*_S_= 1.5 (p=0.584) (Figure 5). We also examined six high-depth samples of pangolin-CoVs from Guangxi, but no conclusions could be drawn due to low sequence quality (Methods; Supplementary Table 18).

To identify recurrent mutations, we next limited to samples for which the major allele was also ancestral at each site (Wuhan-Hu-1 genotype). At such sites, precluding coinfection by multiple genotypes, minor alleles occurring in more than one sample are expected to be derived and recurrent (i.e., identical by state but not descent). For *ORF3d*, we observe *ORF3d*-LOF as a minor allele in two (0.9%) of 220 samples (SRR11410536 and SRR11479046 at frequencies of 1.9% and 4.6%, respectively). This proportion of samples with a recurrent mutation is high but not unusual, as 1.8% of possible genomic changes have an equal or higher proportion of recurrence. Additionally, no mutations in *ORF3d* recur in >2.5% of samples (Figure 8). Thus, we find no evidence that within-host selection or mutation pressure are involved in the spread of *ORF3d*-LOF. However, a small number of other loci exhibit high rates of recurrent mutation, with five mutations independently observed in >10% of samples (Methods; Figure 8). Surprisingly, another STOP mutation (A404U; NSP1-L47*) is observed at low frequency in the majority of samples (Figure 8, Figure 8—figure supplement 1), unexplainable by mutational bias and warranting investigation.

**Figure 8.**
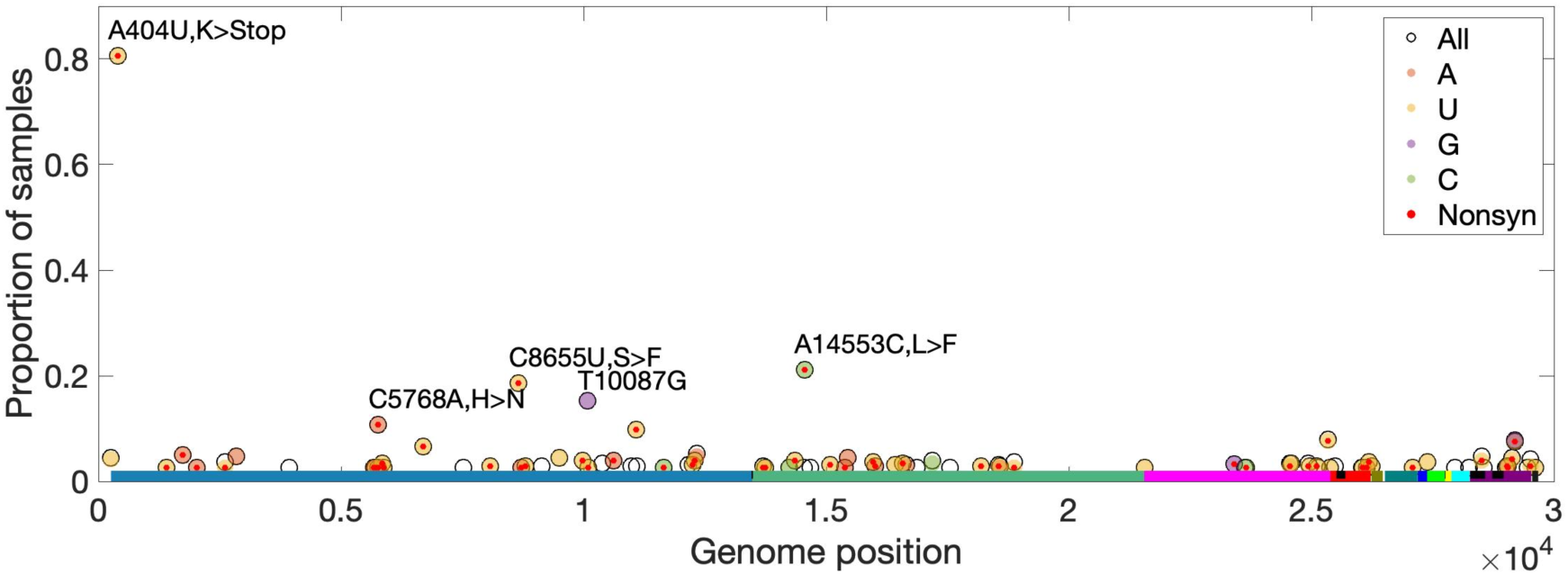
High-frequency within-host mutations. For each site, mutations that occur in more than 2.5% of samples are shown, limiting to samples where the major allele matches Wuhan-Hu-1 (differs by site). The y-axis shows the proportion of such samples having the indicated minor (derived) allele. Each locus has up to three possible single-nucleotide derived alleles compared to the reference background. Open circles (black outlines) show the proportion of samples having any of the three possible derived alleles (“All”), while solid circles (color fill) show the proportion of samples having a specific derived allele (equivalent if only one variant is observed). For most sites, only one derived mutation type (e.g., C→U) was observed across all samples. Precluding co-infection by multiple genotypes and sequencing errors, derived mutations occurring in more than one sample (y-axis) must be identical by state but not descent (i.e., recurrent). Genome positions are plotted on the x-axis, with distinct genes shown in different colors and overlapping genes shown as black blocks within reference genes. Nonsynonymous and nonsense mutations (“Nonsyn”) are indicated with a red dot. Source data available in supplementary material.

## Discussion

Our analyses provide strong evidence that SARS-CoV-2 contains a novel overlapping gene (OLG), *ORF3d*, that has not been consistently identified or fully analyzed before this study. The annotation of a newly emerged virus is difficult, particularly in genome regions subject to frequent gene gain and loss. Moreover, due to the inherent difficulty of studying OLGs, they tend to be less carefully documented than non-OLGs; for example, *ORF9b* and *ORF9c* are still not annotated in the most-used reference genome, Wuhan-Hu-1 (last accessed September 26, 2020). Therefore, *de novo* and homology-based annotation are both essential, followed by careful expression analyses using multi-omic data and evolutionary analyses within and between species. In particular, we emphasize the importance of using whole-gene or genome alignments when inferring homology for both OLGs and non-OLGs, taking into account genome positions and all reading frames. Unfortunately, in the case of SARS-CoV-2, the lack of such inspections has led to mis-annotation and a domino effect. For example, homology between *ORF3b* (SARS-CoV) and *ORF3d* (SARS-CoV-2) has been implied, leading to unwarranted inferences of shared functionality and subsequent claims of homology between *ORF3b* and other putative OLGs within *ORF3a* (Table 1). Given the rapid growth of the SARS-CoV-2 literature, it is likely this mistake will be further propagated. We therefore provide a detailed annotation of Wuhan-Hu-1 protein-coding genes and codons in Supplementary Tables 2 and 3, respectively, as a resource for future studies.

Our study highlights the highly dynamic process of frequent losses and gains of accessory genes in the species *Severe acute respiratory syndrome-related coronavirus*, with the greatest functional constraint typically observed for the most highly expressed genes (Figure 5—figure supplement 1). With respect to gene loss, while many accessory genes may be dispensable for viruses in cell culture, they often play an important role in natural hosts (Forni et al. 2017). Thus, their loss may represent a key step in adaptation to new hosts after crossing a species barrier (Gorbalenya et al. 2006). For example, the absence of full-length *ORF3b* in SARS-CoV-2 has received attention from only a few authors (e.g., Lokugamage et al. 2020), even though it plays a central role in SARS-CoV infection and early immune interactions as an interferon antagonist (Kopecky-Bromberg et al. 2007), with effects modulated by ORF length (Zhou et al. 2012). Thus, the absence or truncation of *ORF3b* in SARS-CoV-2 may be immunologically important (Yuen et al. 2020), e.g., in the suppression of type I interferon induction (Konno et al. 2020). With respect to gene gain, the apparent presence of the *ORF3d* coincident with the inferred entry of SARS-CoV-2 into humans from a hitherto undetermined reservoir host suggests that this gene is functionally relevant for the emergent properties of SARS-CoV-2, analogous to *asp* for HIV-1-M (Cassan et al. 2016). The mechanisms and detailed histories of gene birth, neo-functionalization, and survival of novel genes remain unclear. However, our findings that ORF3d remarkably inherited its hydrophobicity profile from the overlapping region of ORF3a, that ORF3d is depleted for predicted CD8+ and CD4+ T cell epitopes, and that *ORF3d* exhibits potential purifying selection between SARS-CoV-2 and pangolin-CoV GX/P5L, provide strong leads for further research. Indeed, both the structural and immunogenic properties of new viral proteins deserve further attention and are important parts of their evolutionary story.

Failure to account for OLGs can also lead to erroneous inference of (or failure to detect) natural selection in known genes, and *vice versa*. For example, a synonymous variant in one reading frame is very likely to be nonsynonymous in a second overlapping frame. As a result, purifying selection in the second frame usually lowers *d*_S_ (raise *d*_N_/*d*_S_) in the first frame, increasing the likelihood of mis-inferring positive selection (Holmes et al. 2006; Sabath et al. 2008; Nelson et al. 2020). Such errors could, in turn, lead to mischaracterization of the genetic contributions of OLG loci to important viral properties such as incidence and persistence. One potential consequence is misguided countermeasure efforts, e.g., through failure to detect functionally conserved or immunologically important regions.

Although our study focuses on *ORF3d*, other OLGs have been proposed in SARS-CoV-2 and warrant investigation. *ORF3d-2*, a shorter isoform of *ORF3d* (Table 1), shows evidence of higher overall expression than *ORF3d* (Figure 2A). However, analysis of full-length *ORF3d* is confounded by its 20 codon triple overlap with both *ORF3a* and *ORF3c*, making expression and selection results difficult to compare between the two isoforms. Although not evolutionarily novel, the recently discovered *ORF3c* (Table 1) shows deep conservation among viruses of this species (Cagliana et al. 2020; Firth 2020; Jungreis et al. 2020), the lowest *π*_N_/*π*_S_ ratio observed in our between-host analysis (Figure 5), strong evidence for translation in SARS-CoV-2 (Figure 2), and the highest predicted CD8+ T cell epitope density (Figure 3). Within *S*, *S-iORF2* (genome positions 21768-21863) also shows evidence of translation from ribosome profiling (Finkel et al. 2020), and between-host comparisons suggest purifying selection (*π*_N_/*π*_S_=0.22, p=0.0278) (Table 1). We therefore suggest that the SARS-CoV-2 genome could contain additional undocumented or new OLGs.

Our comprehensive evolutionary analysis of the SARS-CoV-2 genome demonstrates that many genes are under relaxed purifying selection, consistent with the exponential growth of the virus (Gazave et al. 2013). At the between-host level, nucleotide diversity increases somewhat over the initial period of the COVID-19 pandemic, tracking the number of locations sampled, while the *π*_N_/*π*_S_ ratio remains relatively constant at 0.46 (±0.030 SEM) (Figure 5—figure supplement 2B). Other genes differ in the strength and direction of selection at the three evolutionary levels, e.g. *ORF9c* (Figure 5), suggesting a shift in function or importance over time or between different host species or individuals. *ORF3d* and *ORF8* are both among the youngest genes in SARS-CoV-2, being taxonomically restricted to a subset of betacoronaviruses (Cui et al. 2019), and both exhibit high rates of change and turnover (Figure 1; Figure 5; SARS-related-CoV_ALN.fasta, supplementary data). High between-host *π*_N_/*π*_S_ was also observed in *ORF8* of SARS-CoV, perhaps due to a relaxation of purifying selection upon entry into civet cats or humans (Forni et al. 2017). However, *ORF3d* and *ORF8* both exhibit strong antibody (B-cell epitope) responses (Hachim et al. 2020) and predicted T-cell epitope depletion (Figure 3) in SARS-CoV-2. This highlights the important connection between evolutionary and immunologic processes (Daugherty and Malik 2012), as antigenic peptides may impose a fitness cost for the virus by allowing immune detection. Thus, the loss or truncation of these genes may share an immunological basis and deserves further attention.

The quick expansion of *ORF3d*-LOF (EP+1+LOF) and its haplotype (EP+1) during this pandemic is surprising, given that a founder effect would have favored other variants that arrived earlier. However, newer data show that *ORF3d*-LOF has remained at relatively low frequencies compared to EP+1 variants (Kosakovsky-Pond 2020), and plausible explanations exist which do not require invoking natural selection. First, the slower ‘spread’ of EP–3 and EP may be an artifact of a sampling policy bias in case isolation: testing and quarantine were preferentially applied to travellers who had recently visited Wuhan, which may have led to selective detection, isolation, and tracing of these haplotypes rather than mutations occurring within Europe (e.g., C14408U and G25563U). However, the EP+1 and EP+1+LOF haplotypes also increase faster in the late founder group, where it is unclear which haplotype was more travel-related. Second, it is possible that the EP haplotype was effectively controlled, while EP+1 was introduced independently (Worobey et al. 2020). Third, although G25563U itself simultaneously causes ORF3a-Q57H and *ORF3d*-LOF, its spread seems to be linked with RdRp-P323L, and both mutations are linked with Spike-D614G, a variant with predicted functional relevance (Korber et al. 2020; Yurkovetskiy et al. 2020). Thus, it is possible that G25563U is neutral while a linked variant(s) is under selection. These observations highlight the necessity of empirically evaluating the effects of these mutations and their interactions. Lastly, because only five major polymorphisms are considered in this analysis (Supplementary Table 14), it is possible that the spread of *ORF3d*-LOF or other haplotypes was further assisted or hindered by subsequent mutations.

Our study has several limitations. First, we were not able to confirm the translation of ORF3d using mass spectrometry (MS). This may be due to several reasons: (1) ORF3d is short, and tryptic digestion of ORF3d generates only two peptides potentially detectable by MS; (2) ORF3d may be expressed at low levels; (3) the tryptic peptides derived from ORF3d may not be amenable to detection by MS even under the best possible conditions, as suggested by its relatively low MS intensity even in an overexpression experiment (‘ORF3b’ in Gordon et al. 2020); and (4) hitherto unknown post-translational modifications of ORF3d could also prohibit detection. Other possibilities for validation of ORF3d include MS with other virus samples; affinity purification MS; fluorescent tagging and cell imaging; Western blotting; and the sequencing of additional genomes in this viral species, which would potentiate more powerful tests of purifying selection and a better understanding of the history and origin of *ORF3d*. With respect to between-host diversity, we focused on consensus-level sequence data; however, this approach can miss important variation (Holmes 2009), stressing the importance of deeply sequenced within-host samples using technology appropriate for calling within-host variants (Grubaugh et al. 2019). As we use Wuhan-Hu-1 for reference-based read mapping and remove duplicate reads as possible PCR artifacts, reference bias (Degner et al. 2009) or bias against natural duplicates at high read depth loci (Zhou et al. 2014) could potentially affect our within-host results. Additionally, we detected natural selection using counting methods that examine all pairwise comparisons between or within specific groups of sequences, which may have less power than methods that trace changes over a phylogeny. However, this approach is robust to errors in phylogenetic and ancestral sequence reconstruction, and to artifacts due to linkage or recombination (Hughes et al. 2006; Nelson and Hughes 2015). Additionally, although our method for measuring selection in OLGs does not explicitly account for mutation bias, benchmarking with other viruses suggests inference of purifying selection is conservative (Nelson et al. 2020). Finally, given multiple recombination breakpoints in *ORF3a* (Boni et al. 2020) and the relative paucity of sequence data for viruses closely related to SARS-CoV-2, our analysis could not differentiate between convergence, recombination, or recurrent loss in the origin of *ORF3d*.

In conclusion, OLGs are an important part of viral biology and deserve more attention. We document several lines of evidence for the expression and functionality of a novel OLG in SARS-CoV-2, *ORF3d*, and compare it to other hypothesized OLG candidates in *ORF3a*. Finally, as a resource for future studies, we provide a detailed annotation of the SARS-CoV-2 genome and highlight mutations of potential relevance to the within- and between-host evolution of SARS-CoV-2.

## Methods

### Data and software

Supplementary scripts and documentation are freely available on GitHub at https://github.com/chasewnelson/SARS-CoV-2-ORF3d. Supplementary data not included in main figure source data are freely available on Zenodo at https://zenodo.org/record/4052729.

### Genomic features and coordinates

All genome coordinates are given with respect to the Wuhan-Hu-1 reference sequence (NCBI: NC_045512.2; GISAID: EPI_ISL_402125) unless otherwise noted. SARS-CoV (SARS-CoV-1) genome coordinates are given with respect to the Tor2 reference sequence (NC_004718.3). SARS-CoV-2 Uniprot peptides were obtained from https://viralzone.expasy.org/8996, where *ORF9c* is referred to as *ORF14*. Nucleotide sequences were translated using R::Biostrings (Lawrence et al. 2013), Biopython (Cock et al. 2009), or SNPGenie (Nelson et al. 2015). Alignments were viewed and edited in AliView v1.20 (Larsson 2014). To identify OLGs using the codon permutation method of Schlub et al. (2018), all 12 ORFs annotated in the Wuhan-Hu-1 reference genome were used as a reference (NCBI=NC_045512.2; *ORF1a*, *ORF1b*, *S*, *ORF3a*, *E*, *M*, *ORF6*, *ORF7a*, *ORF7b*, *ORF8*, *N*, and *ORF10*) (Figure 1; Supplementary Tables 1-3). Only forward-strand (same-strand; sense-sense) OLGs were considered in this analysis.

### SARS-CoV-2 genome data, alignments, and between-host analyses

SARS-CoV-2 genome sequences were obtained from GISAID on April 10, 2020 (GISAID Acknowledgments Table, supplementary data). Whole genomes were aligned using MAFFT v7.455 (Katoh and Standley 2013), and subsequently discarded if they contained internal gaps (-) located >900 nt from either terminus, a length sufficient to exclude sequences with insertions or deletions (indels) in protein-coding regions. A total of 3,978 sequences passed these filtering criteria, listed in Supplementary Table 13. Coding regions were identified using exact or partial homology to SARS-CoV-2 or SARS-CoV annotations. To quantify the diversity and evenness of sample locations, we quantified their entropy as -∑*p*\*ln(*p*), where *p* is the number of distinct (unique) locations or countries reported for a given window (Ewens and Grant 2001) (Supplementary Table 19).

### Severe acute respiratory syndrome-related coronavirus genome data and alignments

*Severe acute respiratory syndrome-related coronavirus* genome IDs were obtained from Lam et al. (2020) and downloaded from GenBank or GISAID. Genotype Wuhan-Hu-1 was used to represent SARS-CoV-2. Except for pangolin-specific analyses, isolates GX/P5L (NCBI: MT040335.1; GISAID: EPI_ISL_410540) and GD/1 (GISAID: EPI_ISL_410721) were used to represent pangolin-COVs; GD/1 was chosen as a representative because the other Guangdong (GD) sequence lacks the *S* gene and contains 27.76% Ns, while GX/P5L was chosen because it is one of two high-coverage Guangxi (GX) sequences derived from lung tissue that also contains no Ns. Other sequences were excluded if they lacked an annotated *ORF1ab* with a ribosomal slippage, or contained a frameshift indel in any gene (Supplementary Table 6), leaving 21 sequences for analysis (Supplementary Table 5).

To produce whole-genome alignments, we first aligned all genome sequences using MAFFT. Then, coding regions were identified using exact or partial sequence identity to SARS-CoV-2 or SARS-CoV annotations, translated, and individually aligned at the amino acid level using ProbCons v1.12 (Do et al. 2005). In the case of OLGs, the longest (reference) protein was used (e.g., N rather than ORF9b or ORF9c). Amino acid alignments were then imposed on the coding sequence of each gene using PAL2NAL v14 (Suyama et al. 2006) to maintain intact codons. Finally, whole genomes were manually shifted to match the individual codon alignments in AliView. Codon breaks were preferentially resolved to align S/Q/T at 3337-3339, and L/T/I at 3343-3345. This preserved all nucleotides of each genome while concurrently producing codon-aware alignments. The alignment is available in the supplementary data as SARS-related-CoV_ALN.fasta, where pangolin-CoV GD/1 has been removed because it is only available from GISAID (i.e., permission is required for data access).

Phylogenetic relationships among isolates were explored using maximum likelihood phylogenetic inference, as implemented in IQ-tree (Nguyen et al. 2015), using the generalized time-reversible (GTR; Tavaré 1986) substitution model combined with the FreeRate model (Soubrier et al. 2012) to account for among-site rate heterogeneity. Trees were rooted *a priori* following Lam et al. (2020).

### Proteomics analysis

We used MaxQuant (Tyanova et al. 2016) to re-analyze five publicly available SARS-CoV-2 mass spectrometry (MS) datasets: Bezstarosti et al. 2020 (PRIDE accession PXD018760); Bojkova et al. 2020 (PXD017710); Davidson et al. 2020 (PXD018241); PRIDE Project PXD018581; and Zecha et al. 2020 (PXD019645). However, peptide spectrum matches for ORF3d (possible tryptic peptides CTSCCFSER and FQNHNPQK) did not pass our 1% false-discovery threshold in any of these datasets, and ORF3d-2 (Table 1) does not encode any peptides detectable by this method. ORF3c, ORF9c, and ORF10 could also not be reliably detected. For peptides that were successfully detected in the datasets of Bezstarosti et al. (2020) and Davidson et al. (2020), protein concentrations were estimated using intensity-based absolute quantification (iBAQ) (Schwanhäusser et al. 2011). iBAQ values were computed using the Max-Quant software (Cox and Mann 2008) as the sum of all peptide intensities per protein divided by the number of theoretical peptides per protein. Thus, iBAQ values are proportional estimates of the molar protein quantity of a protein in a given sample, allowing relative quantitative comparisons of protein expression levels.

### Ribosome profiling analysis

The 16 ribosome profiling (Ribo-seq) datasets of Finkel et al. (2020) using SARS-CoV-2 infected Vero E6 cells were downloaded from the Sequence Read Archive (accession numbers SRR11713354 to SRR11713369). These samples comprise four treatments for ribosome stalling: (1) lactimidomycin (LTM) or (2) harringtonine (Harr), which are biased towards stalling at translation initiation sites; (3) cyclohexamide (CHX), which stalls along a gene during active translation; and (4) mRNA, which serves as a control (i.e., total RNA content rather than ribosome footprints). Two time points are represented: (1) 5 hours and (2) 24 hours post infection (hpi), labelled “05hr” and “4hr”, respectively, in the SRA data. The FASTQ format reads were mapped to the Wuhan-Hu-1 reference genome using Bowtie2 local alignment (Langmead et al. 2019), with a seed length of 20 and up to one mismatch allowed, after substituting the isolate’s mutations, as listed in Finkel et al. (2020). Mapped reads were then classified by read length and trimmed to the first (5′-end) nucleotide for downstream analyses.

### NetMHCpan T-cell epitope analysis

For predicted MHC class I binding, viral protein sequences were analyzed in 9 aa substrings using NetMHCpan4.0 (Jurtz et al. 2017). Twelve (12) representative HLA class I alleles (Sidney et al. 2008) were tested: HLA-A*01:01 (A1), HLA-A*02:01 (A2), HLA-A*03:01 (A3), HLA-A*24:02 (A24), HLA-A*26:01 (A26), HLA-B*07:02 (B7), HLA-B*08:01 (B8), HLA-B*27:05 (B27), HLA-B*39:01 (B39), HLA-B*40:01 (B44), HLA-B*58:01 (B58), and HLA-B*15:01 (B62). NetMHCpan4.0 returns percentile ranks that characterize a peptide’s likelihood of antigen presentation compared to a set of random natural peptides. We employed the suggested threshold of 2% to determine potential presented peptides, and 0.5% to identify strong MHC class I binder. Both strong and weak binders were considered predicted epitopes. For predicted MHC class II binding, viral protein sequences were analyzed in 15 aa substrings using NetMHCIIpan4.0 (Reynission et al. 2020). Twenty six (26) representative HLA class II alleles (Greenbaum et al. 2011; Paul et al. 2016) were tested: HLA-DPA1*0201-DPB1*0401, HLA-DPA1*0103-DPB1*0201, HLA-DPA1*0201-DPB1*0101, HLA-DPA1*0201-DPB1*0501, HLA-DPA1*0301-DPB1*0402, HLA-DQA1*0101-DQB1*0501, HLA-DQA1*0102-DQB1*0602, HLA-DQA1*0301-DQB1*0302, HLA-DQA1*0401-DQB1*0402, HLA-DQA1*0501-DQB1*0201, HLA-DQA1*0501-DQB1*0301, DRB1*0101, DRB1*0301, DRB1*0401, DRB1*0405, DRB1*0701, DRB1*0802, DRB1*0901, DRB1*1101, DRB1*1201, DRB1*1302, DRB1*1501, DRB3*0101, DRB3*0202, DRB4*0101, DRB5*0101. NetMHCIIpan4.0 returns percentile ranks that characterize a percentile rank of eluted ligand prediction score. We employed the suggested threshold of 10% to determine potential presented peptides, and 2% to identify strong MHC class II binder. Both strong and weak binders were considered predicted epitopes.

### Hydrophobicity profile analysis

The sequence of *ORF3a* was translated in all three forward-strand reading frames and uploaded to the VOLPES server (http://volpes.univie.ac.at/; Bartonek and Zagrovic 2019) to determine hydrophobicity profiles using the unitless scale “FAC1” (Factor 1; Atchley et al. 2005) with a window size of 25 codons. The correlations between the profiles for the reading frames were calculated for peptides encoded by the following subregions of *ORF3a*, from 5′ to 3′: (1) *ORF3a*, codons not overlapping any known or hypothesized overlapping genes; (2) *ORF3a*/*ORF3c*, encoding residues of both *ORF3a* and *ORF3c*; (3) *ORF3a*/*ORF3c*/*ORF3d*, encoding residues of *ORF3a*, *ORF3c*, and *ORF3d*; (4) *ORF3a*/*ORF3d*, encoding residues of both *ORF3a* and *ORF3d*; and (5) *ORF3a*/*ORF3b*, encoding residues of both *ORF3a* and *ORF3b*. Spearman’s rank correlation was computed using stats::cor.test() with method=’spearman’ in R, treating mean 25-residue window values as the observational unit.

### Statistics and tests of natural selection

All p-values reported in this study are two-tailed. Statistical and data analyses and visualization were carried out in R v3.5.2 (R Core Team 2018) (libraries: boot, feather, ggrepel, patchwork, RColorBrewer, scales, tidyverse), Python (BioPython, pandas) (McKinney 2010), Microsoft Excel, Google Sheets, and PowerPoint. Colors were explored using Coolors (https://coolors.co). Copyright-free images of a bat, human, and pangolin were obtained from Pixabay (https://pixabay.com). All *d*_N_/*d*_S_ or *π*_N_/*π*_S_ ratios were estimated for non-OLG regions using SNPGenie scripts snpgenie.pl or snpgenie_within_group.pl (Nelson et al. 2015; https://github.com/chasewnelson/SNPGenie), and for OLG regions using OLGenie script OLGenie.pl (Nelson et al. 2020; https://github.com/chasewnelson/OLGenie). All OLG *d*_N_/*d*_S_ (*π*_N_/*π*_S_) estimates refer to *d*_NN_/*d*_SN_ (*π*_NN_/*π*_SN_) for the reference frame and *d*_NN_/*d*_NS_ (*π*_NN_/*π*_NS_) for the alternate frame (ss12 or ss13), as described in Nelson et al. (2020), because the number of SS (synonymous/synonymous) sites was insufficient to estimate *d*_SS_ (*π*_SS_). The null hypothesis that *d*_N_-*d*_S_=0 (*π*_N_-*π*_S_=0) was evaluated using both *Z* and achieved significance level (ASL) tests (Nei and Kumar 2000) with 10,000 and 1,000 bootstrap replicates for genes and sliding windows, respectively, using individual codons (alignment columns) as the resampling unit (Nei and Kumar 2000). The main text reports only *Z*-test results, because this test was used to benchmark OLGenie (Nelson et al. 2020) and appears to be more conservative. For ASL, p-values of 0 were reported as the lowest non-zero value possible given the number of bootstrap replicates. Benjamini-Hochberg (Benjamini and Hochberg 1995) or Benjamini-Yekutieli (Benjamini and Yekutieli 2001) false-discovery rate corrections (*Q*-values) were used for genes (independent regions) and sliding windows (contiguous overlapping regions), respectively.

### Between-species analyses

Because uncorrected *d* values >0.1 were observed in between-taxa comparisons, a Jukes-Cantor correction (Jukes and Cantor 1969) was applied to *d*_N_ and *d*_S_ estimates. For each ORF, sequences were only used to estimate *d*_N_/*d*_S_ if a complete, intact ORF (no STOPs) was present. Most notably, for *ORF3b*, only the region corresponding to the first ORF in SARS-CoV-2 was analyzed, and these sites were also considered an OLG region of *ORF3a*. The remaining 3’-proximal portion of *ORF3a* (codons 165-276) was considered a non-OLG region, and only genomes lacking the full-length *ORF3b* (i.e., those with mid-sequence STOP codons) were included in these estimates. Similarly, for *ORF3d*, only SARS-CoV-2 and pangolin-CoV GX/P5L were analyzed for Figure 5. For this analysis, *ORF3a* codon 71 in SARS-CoV-2 (CTA) differed by two nucleotides from the same codon in pangolin-CoV GX/P5L (TTT), resulting in a multi-hit approximation being employed by the OLGenie method (Nelson et al. 2020). Thus, for greater accuracy, we instead estimated changes in this codon using the method of Wei and Zhang (2015) across two mutational pathways, which yields 0.5 nonsynonymous/nonsynonymous, 0.5 nonsynonymous/synonymous, and 1 synonymous/nonsynonymous changes. The region occupied by the triple overlap of *ORF3a*/*ORF3c*/*ORF3d* (*ORF3a* codons 44-64; Table 1) was excluded from analysis for all three genes. The following codons (or their homologous positions) were also excluded from all analyses: all codons occupying a non-OLG/OLG boundary; codons 4460-4466 of *ORF1ab*, which constitute either *nsp11* or *nsp12* depending on a ribosomal frameshift; codons 1-13 of *E*, which overlap *ORF3b* in some genomes; codons 62-64 of *ORF6*, which follow a premature STOP in some genomes; codons 122-124 of *ORF7a* and 1-3 of *ORF7b*, which overlap each other; and codons 72-74 of *ORF9c*, which follow a premature STOP in some genomes.

### Cumulative haplotype frequencies

We defined haplotypes along the mutational path to EP+1+LOF using all five high-derived allele frequency (DAF) mutations from Wuhan-Hu-1 to *ORF3d*-LOF. Subsequent mutations after *ORF3d*- LOF were ignored in the haplotype analysis. Samples with missing data at any of the five loci were removed from the haplotype analysis. We calculated the cumulative frequency of each haplotype in Germany (where the EP haplotype is a documented founder) and five other countries with the most abundant samples by the time of data accession. Cumulative frequencies were calculated as the total number of occurrences of each haplotype collected each day, divided by the total number of samples from the same country. Countries were subsequently divided into early founders and late founders to investigate founder effects, where early founder countries tend to have several samples from January, and late founder countries tend to have samples collected only after mid-February.

### Within-host diversity

For within-host analyses, we obtained *n*=401 high-depth (at least 50-fold mean coverage) human SARS-CoV-2 samples from the Sequence Read Archive (listed in Supplementary Table 17). Only Illumina samples were used, as some Nanopore samples exhibit apparent systematic bias in calling putative intrahost SNPs, and this technology has also been shown to be unsuitable for intra-host analysis (Grubaugh et al. 2019). Reads were trimmed with BBDUK (Bushnell B. 2017. BBTools. https://jgi.doe.gov/data-and-tools/bbtools/) and mapped against the Wuhan-Hu-1 reference sequence using Bowtie2 (Langmead and Salzberg 2012) with local alignment, seed length 20, and up to 1 mismatch. SNPs were called from mapped reads using the LoFreq (Wilm et al. 2012) variant caller with sequencing quality and MAPQ both at least 30. Only single-end or the first of paired-end reads were used. Variants were dynamically filtered based on each site’s coverage using a binomial cutoff to ensure a false-discovery rate of ≤1 within-host variant across our entire study (401 samples), assuming a mean sequencing error rate of 0.2% (Schirmer et al. 2016).

To estimate *π*, numbers of nonsynonymous and synonymous differences and sites were first calculated individually for all genes in each of the 401 samples using SNPGenie (Nelson et al. 2015; snpgenie.pl, https://github.com/chasewnelson/SNPGenie). OLG regions were then analyzed separately using OLGenie (Nelson et al. 2020; OLGenie.pl, https://github.com/chasewnelson/OLGenie). Because OLGenie requires a multiple sequence alignment as input, a pseudo-alignment of 1,000 sequences was constructed for each OLG region in each sample by randomly substituting single nucleotide variants according to their within-host frequencies into the Wuhan-Hu-1 reference genome. For non-OLG and OLG regions alike, average within-host numbers of differences and sites were calculated for each codon by taking the mean across all samples. For example, if a particular codon contained nonsynonymous differences in two of 401 samples, with the two samples exhibiting mean numbers of 0.01 and 0.002 pairwise differences per site, this codon was considered to exhibit a mean of (0.01+0.002)/401=0.0000299 pairwise differences per site across all samples. These codon means were then treated as independent units of observation during bootstrapping.

Pangolin samples refer to Sequence Read Archive records SRR11093266, SRR11093267, SRR11093268, SRR11093269, SRR11093270, and SRR11093271. Only 179 single nucleotide variants could be called prior to our FDR filtering, and samples SRR11093271 and SRR11093270 were discarded entirely due to low mapping quality. We also note that after our quality filtering, four samples contain consensus alleles that do not match their reference sequence (available from GISAID): P1E, P4L, P5E, and P5L (Supplementary Table 18).

### Within-host recurrent mutations analyses

We assume that each host was infected by a single genotype. Under this assumption, the minor allele of each segregating site within-host is either due to genotyping or sequencing artifacts, or new mutations. Because there are very few loci with high-frequency derived alleles between hosts, and because the Wuhan-Hu-1 genome is used as the reference in read mapping, we here only consider within-host mutations that differ from this reference background. There are four possible bases at each locus, A, C, G, and U, and three possible mutational changes away from the Wuhan-Hu-1 reference genome. For each locus, we calculate the number of samples matching the reference allele as *N*=*N*_1_+*N*_2_, where *N*_1_ is the number of samples in which the Wuhan reference allele is the only observed allele, and *N*_2_ is the number of samples in which the Wuhan reference allele is major allele. Given *N*_2_, we further determined the number of samples carrying each of the three possible non-reference alleles as *N*_A_, *N*_C_, *N*_G_, and *N*_U_. For example, if the reference allele was U, we calculated *p*_A_, *p*_C_, and *p*_G_, where *p*_All_=*p*_A_+*p*_C_+*p*_G_. If A was an observed non-reference allele, we calculated the frequency of A as *p*_A_=*N*_A_/*N*. Thus, a larger frequency indicates the derived allele is observed in a high proportion of samples (Figure 8). The within-host derived allele frequency (DAF) was estimated as the total number of reads matching the observed minor allele divided by the total number of reads mapped to the locus. If all reads matched the Wuhan-Hu-1 reference allele, then DAF=0. Five mutations occur in more than 10% of samples, four of which are nonsynonymous, with their DAF plotted in Figure 8—figure supplement 1. For this analysis, we did not apply the per-site FDR cutoff, thus a DAF=0 is equivalent to the absence of reads mapped to the mutation, after reads are filtered by sequence quality, mapping quality, and LoFreq’s default significance threshold (p=0.01).

## Supporting information

Supplemental GISAID Acknowledgment Table

## Author Contributions

X.W. conceived the study. C.W.N., Z.A., and X.W. designed the study. T.G., S.-O.K., and X.W., advised on the study. C.W.N. and Z.A. obtained and processed data. C.W.N., Z.A., C-H.K., M.C., C.L., S.-O.K., and X.W., analyzed data. C.W.N., Z.A., S.-O.K., and X.W. conceived, discussed, and illustrated figures. All authors discussed and interpreted results. C.W.N., Z.A., T.G., S.-O.K., and X.W. wrote the first draft. All authors read, discussed, and revised the manuscript.

## Acknowledgements

This work was supported by a Postdoctoral Research Fellowship from Academia Sinica (to C.W.N. under P.I. Wen-Hsiung Li); funding from the Bavarian State Government and National Philanthropic Trust (to Z.A. under P.I. Siegfried Scherer); NSF IOS grants #1755370 and #1758800 (to S.-O.K.); and the University of Wisconsin-Madison John D. MacArthur Professorship Chair (to T.L.G). Copyright-free images were obtained from Pixabay. The authors thank the GISAID platform and the originating and submitting laboratories who kindly uploaded SARS-CoV-2 sequences to the GISAID EpiCov™ Database for public access (see GISAID Acknowledgments Table in supplement). The authors thank Maciej F. Boni, Reed A. Cartwright, John Flynn, Kyle Friend, Dan Graur, Robert S. Harbert, Cheryl Hayashi, David G. Karlin, Niloufar Kavian, Kin-Hang (Raven) Kok, Wen-Hsiung Li, Meiyeh Lu, David A. Matthews, Lisa Mirabello, Apurva Narechania, Felix Li Jin, and attendees of the UC Berkeley popgen journal club for useful information and discussion; Andrew E. Firth, Alexander Gorbalenya, Irwin Jungreis, Manolis Kellis, Raven Kok, Angelo Pavesi, Kei Sato, Manuela Sironi, and Noam Stern-Ginossar for an invaluable discussion regarding standardizing nomenclature; Helen Piontkivska, Patricia Wittkopp, Antonis Rokas, and one anonymous reviewer for critical suggestions; Ming-Hsueh Lin for immense feedback on figures; and special thanks to Priya Moorjani, Jacob Tennessen, Montgomery Slatkin, Yun S. Song, Jianzhi George Zhang, Xueying Li, Hongxiang Zheng, Qinqin Yu, Meredith Yeager, and Michael Dean for commenting on earlier drafts of this manuscript.

---

**Figure 1—figure supplement 1.**
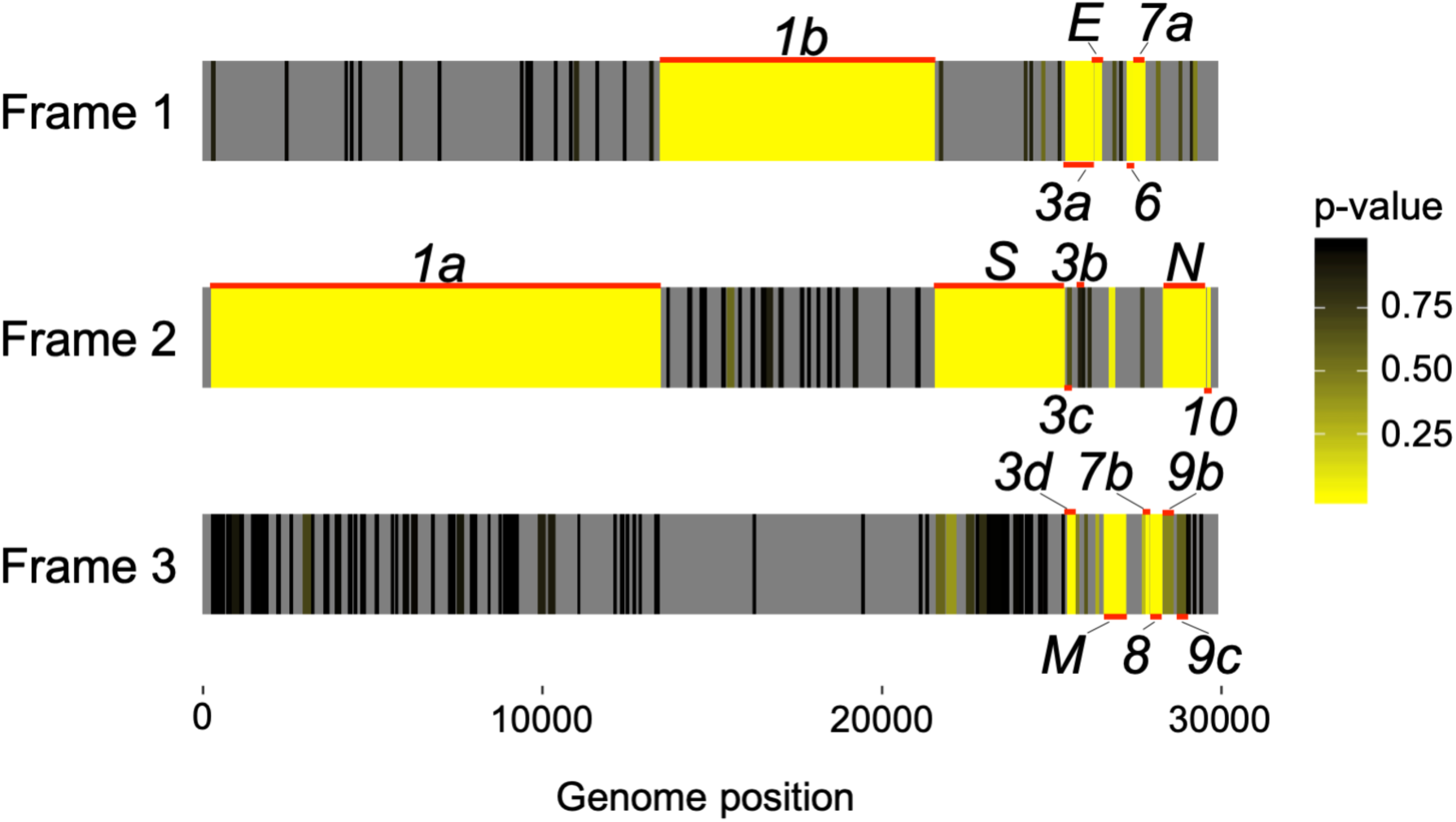
Codon permutation analysis to identify candidate overlapping genes in all three forward-sense reading frames of the SARS-CoV-2 genome. Genome positions of known and hypothesized genes are indicated with horizontal red lines. Reading frames 1, 2, and 3 begin at position 1, 2, or 3 of the Wuhan-Hu-1 reference genome, respectively, with genome coordinates shown at the bottom (Supplementary Tables 2 and 3). Yellow indicates low p-values (natural logarithm scale), while gray indicates absence of a stop-to-stop ORF longer than 30 codons or presence outside of a known canonical gene (not tested). Note that low p-values are expected for the non-overlapping genes. Only the first of four SARS-CoV-2 ORFs homologous to *ORF3b* is shown (Figure 4; Supplementary Table 4).

**Figure 2—figure supplement 1.**
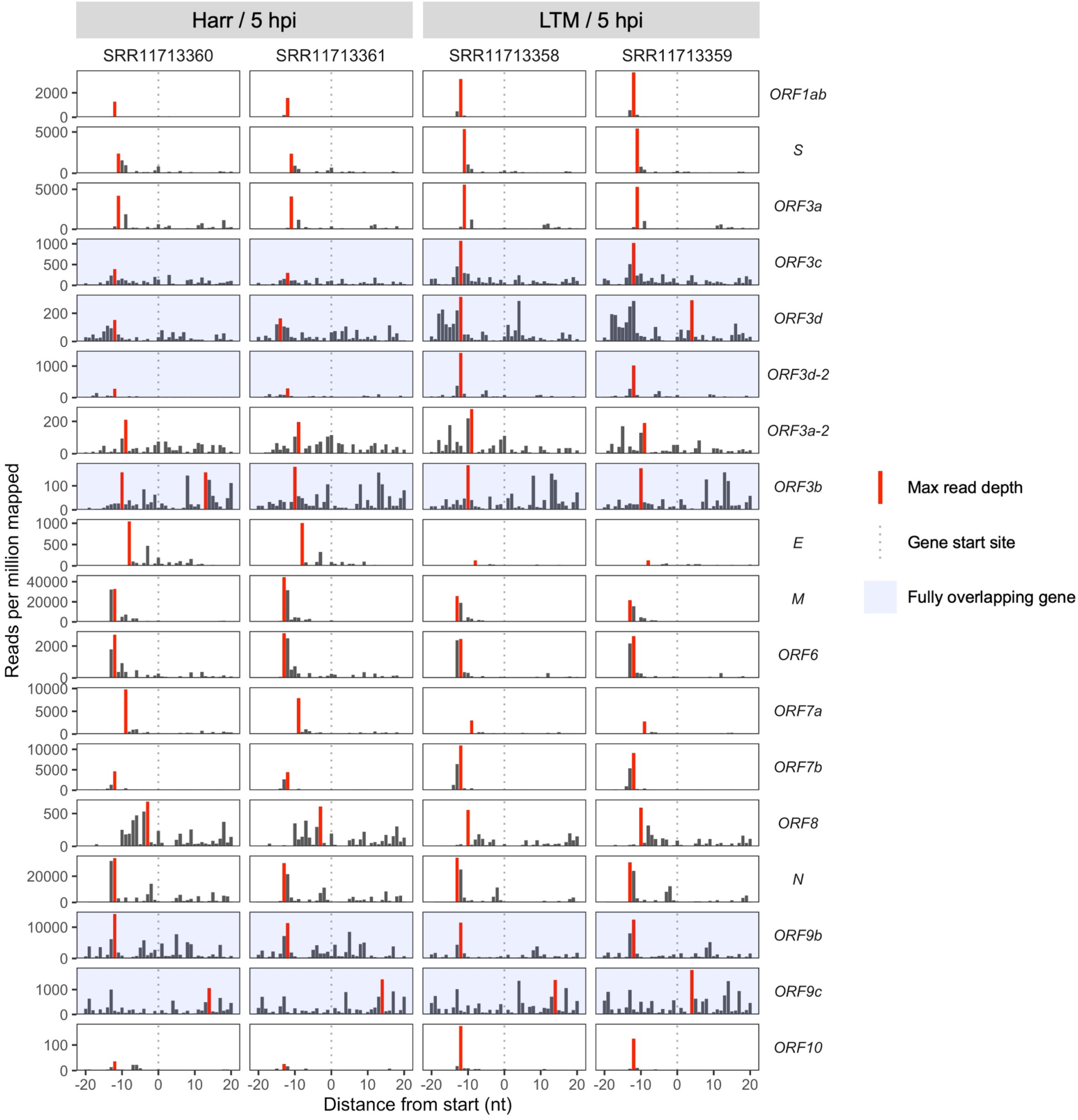
Ribosome profiling coverage near translation initiation sites for individual samples. Details as in Figure 2A, except samples were not pooled and genes are arranged vertically by start site in the genome from 5′ to 3′ (top to bottom). Treatments are indicated as Harr=harringtonine and LTM=lactimidomycin at 5 hours post-infection (hpi).

**Figure 2—figure supplement 2.**
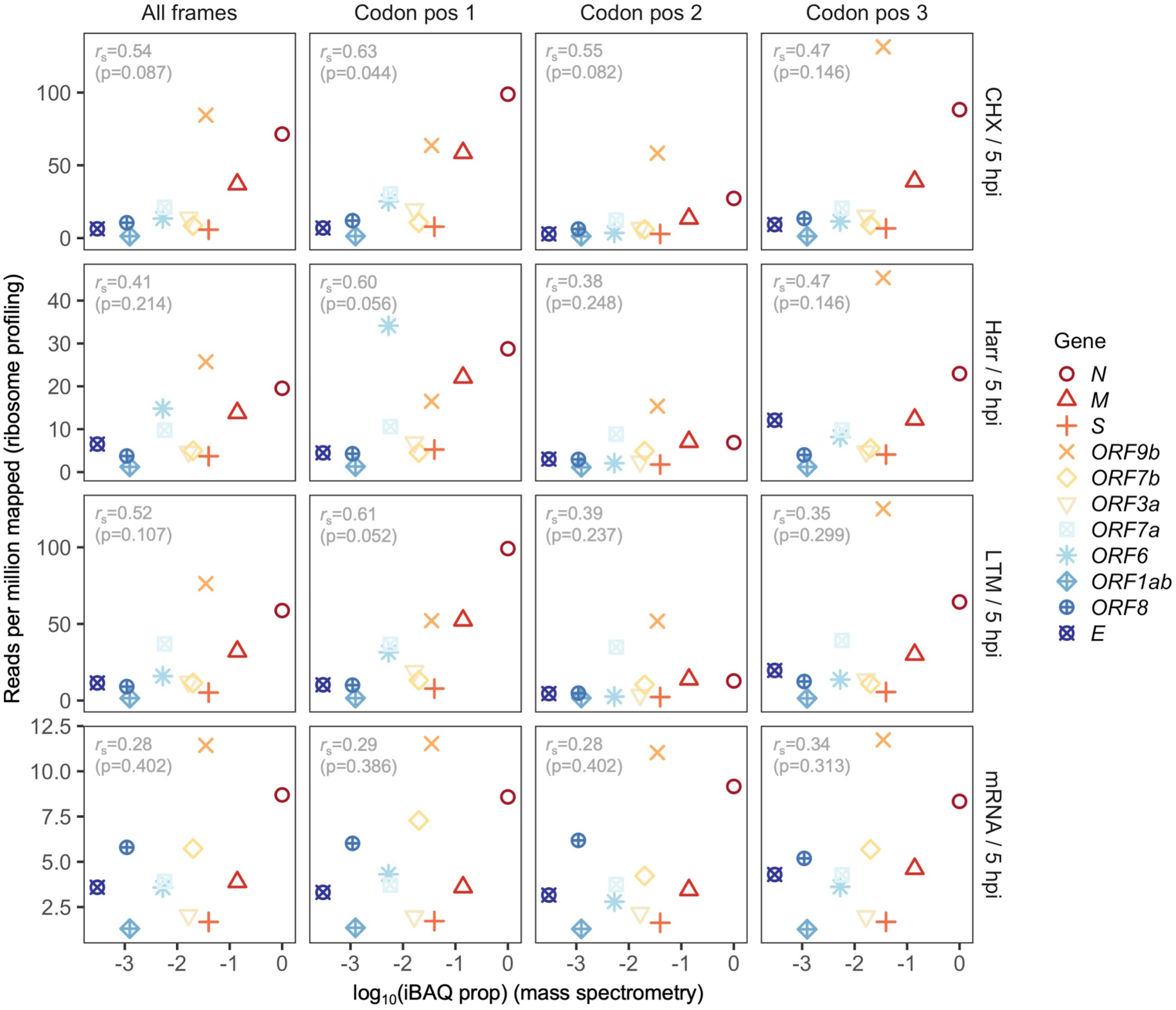
Correlation between protein expression as estimated by mass spectrometry and ribosomal profiling for individual treatments and codon positions. Details as in Figure 2B, except estimates are shown for all sites in a gene (All frames) and out-of-frame reads are shown separately for codon positions (Codon pos) 2 and 3. *ORF9b* is a fully (internal) overlapping gene occurring within *N*, with codon position 3 of *ORF9b* corresponding to codon position 1 of *N*. As a result, codon position 3 of *ORF9b* is predominantly occupied by ribosomes translating *N*, the most highly expressed gene, leading to overestimation of *ORF9b* expression (Codon pos 3; right). Ribosome profiling read depth at codon position 1 exhibits a marginal Spearman’s rank correlation with mass spectrometry for cycloheximide (CHX), harringtonine (Harr), and lactimidomycin (LTM) treatments; however, as expected, all correlations for RNA-seq (mRNA), as well as reads mapping to other codon positions, were not significant.

**Figure 2—figure supplement 3.**
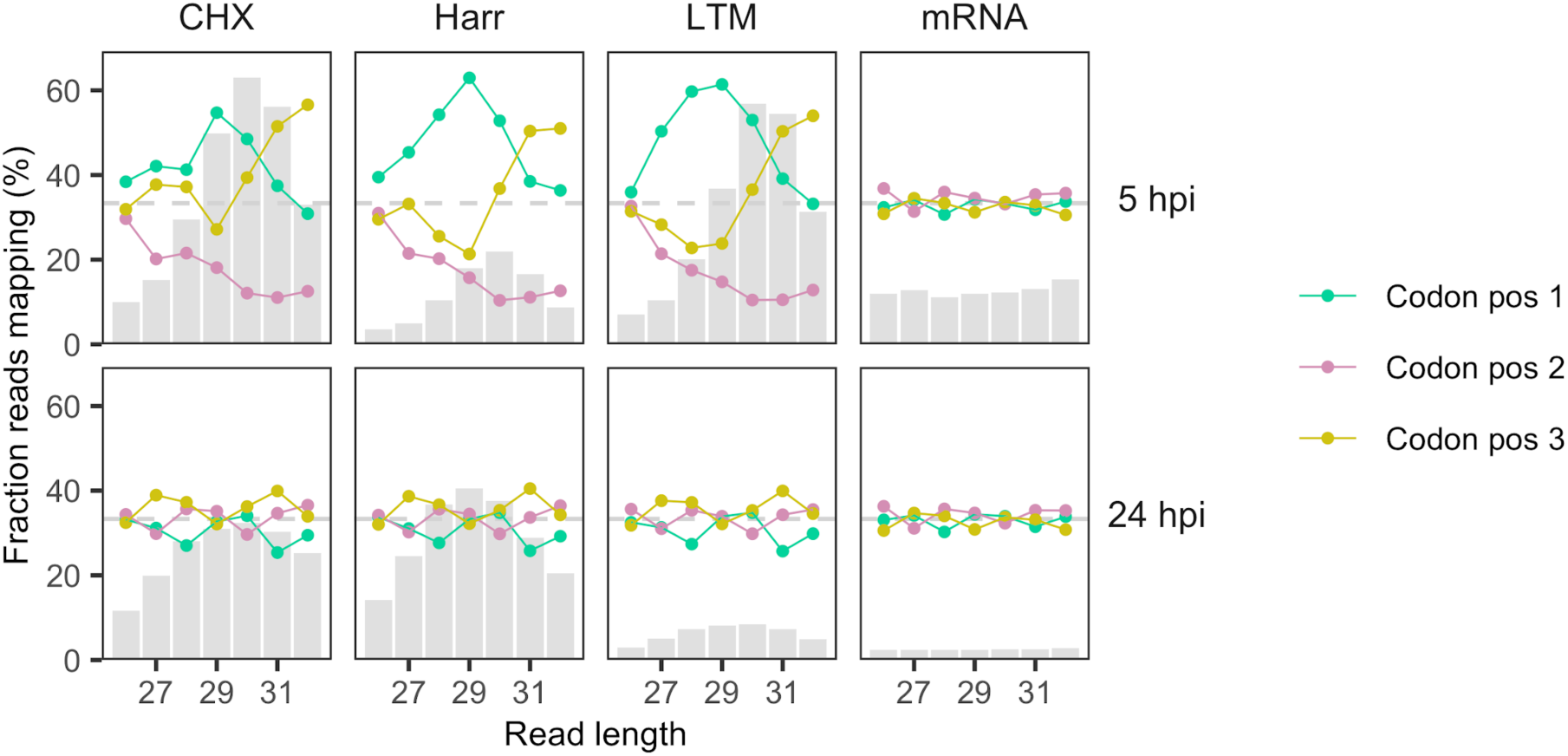
Reading frames occupied by the 5′ ends of ribosome profiling reads as a function of read length. Processed, aligned reads from Finkel et al. (2020) were trimmed to their 5′ ends (first nt). Fractions of reads mapping to each codon position (x axis) are shown as a function of read length (y axis; 26-32 nt) after pooling samples according to treatment (CHX=cycloheximide, Harr=harringtonine, LTM=lactimidomycin, mRNA=RNA-seq) and time (hours post-infection; hpi). Codon position is defined relative to gene start sites of annotated (canonical) non-overlapping genes, excluding any overlapping gene regions. Gray bars show relative total counts for reads of different lengths, with different scales for 5 hpi (lower coverage; max=284,608 reads at CHX/30nt) and 24 hpi (higher coverage; max=3,149,251 reads at Harr/29nt). At 5 hpi, reads of length 29 and 30 nt tend to have 5′ ends mapping to the correct protein-coding reading frame (codon position 1) of the gene in which they occur, and also tend to be most abundant, as expected given the size of ribosome-protected fragments (∼30 nt). Samples from 24 hpi have flatter (left-shifted) read length distributions and no clear frame signal. The horizontal dashed grey line corresponds to the expectation for RNA fragments not protected by ribosomes, i.e., 33% of reads mapping to each codon position (random).

**Figure 2—figure supplement 4.**
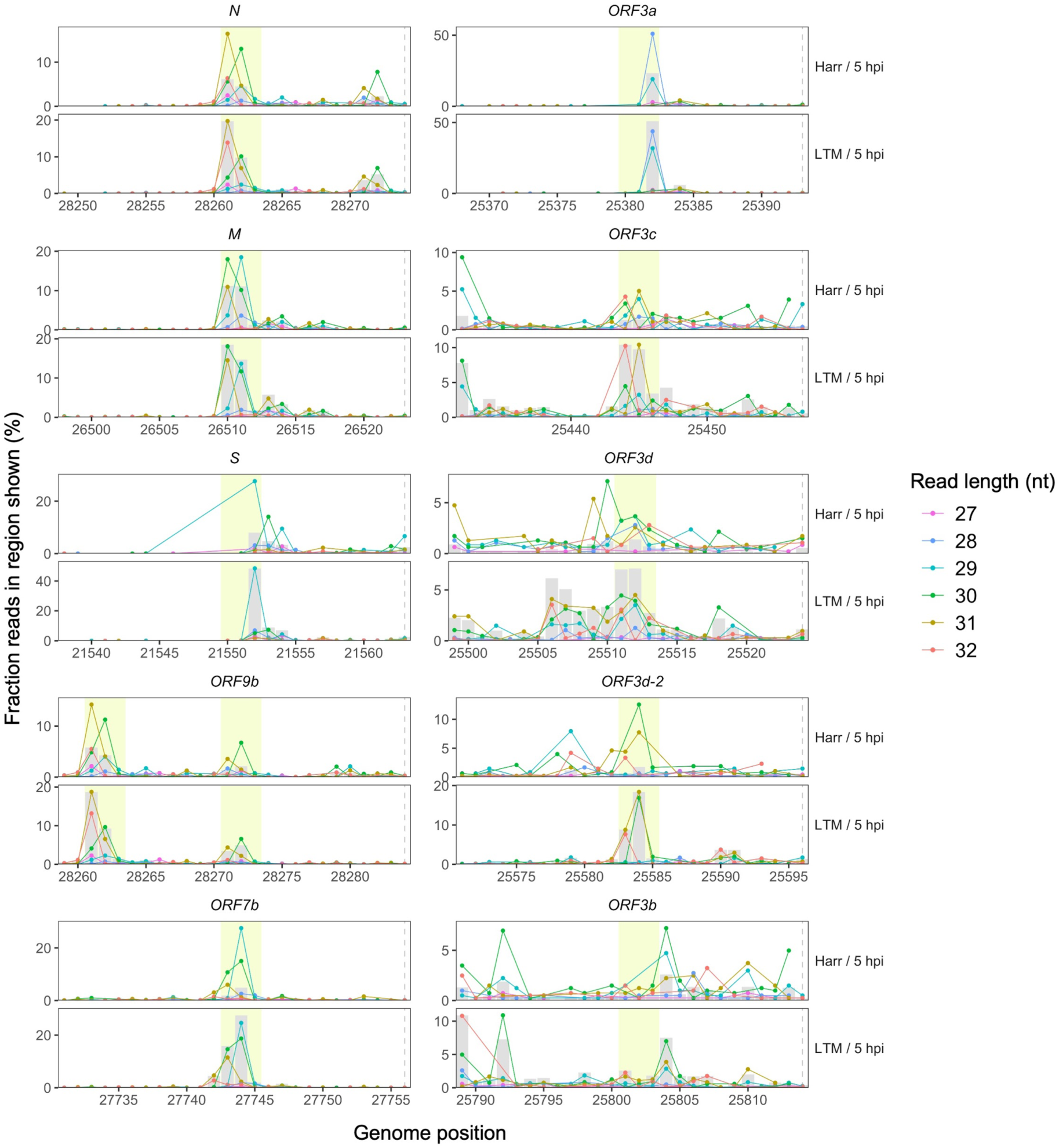
Ribosome profiling read accumulation at gene start sites as a function of read length. Points indicate the percentage of all reads in the region shown whose first nucleotides (5′ ends) map to each genome position, colored according to read length. In each case, the longer read lengths have 5′ ends that predominate at approximately -12 nt from the gene start site (horizontal dashed lines, right), corresponding to the ribosome P-site offset (center of yellow highlight). A peak in total read depth (grey bars) is also observed at this position, as seen in Figure 2A. Highly expressed canonical (*N*, *M*, *S*, and *ORF7b*) and overlapping (*ORF9b*) genes are shown on the left, while *ORF3a* and its hypothesized overlapping genes are shown on the right. Note that *ORF9b* begins shortly after *N*, such that the upstream peak of *N* occurs at position -22 nt relative to the start site of *ORF9b*.

**Figure 2—figure supplement 5.**
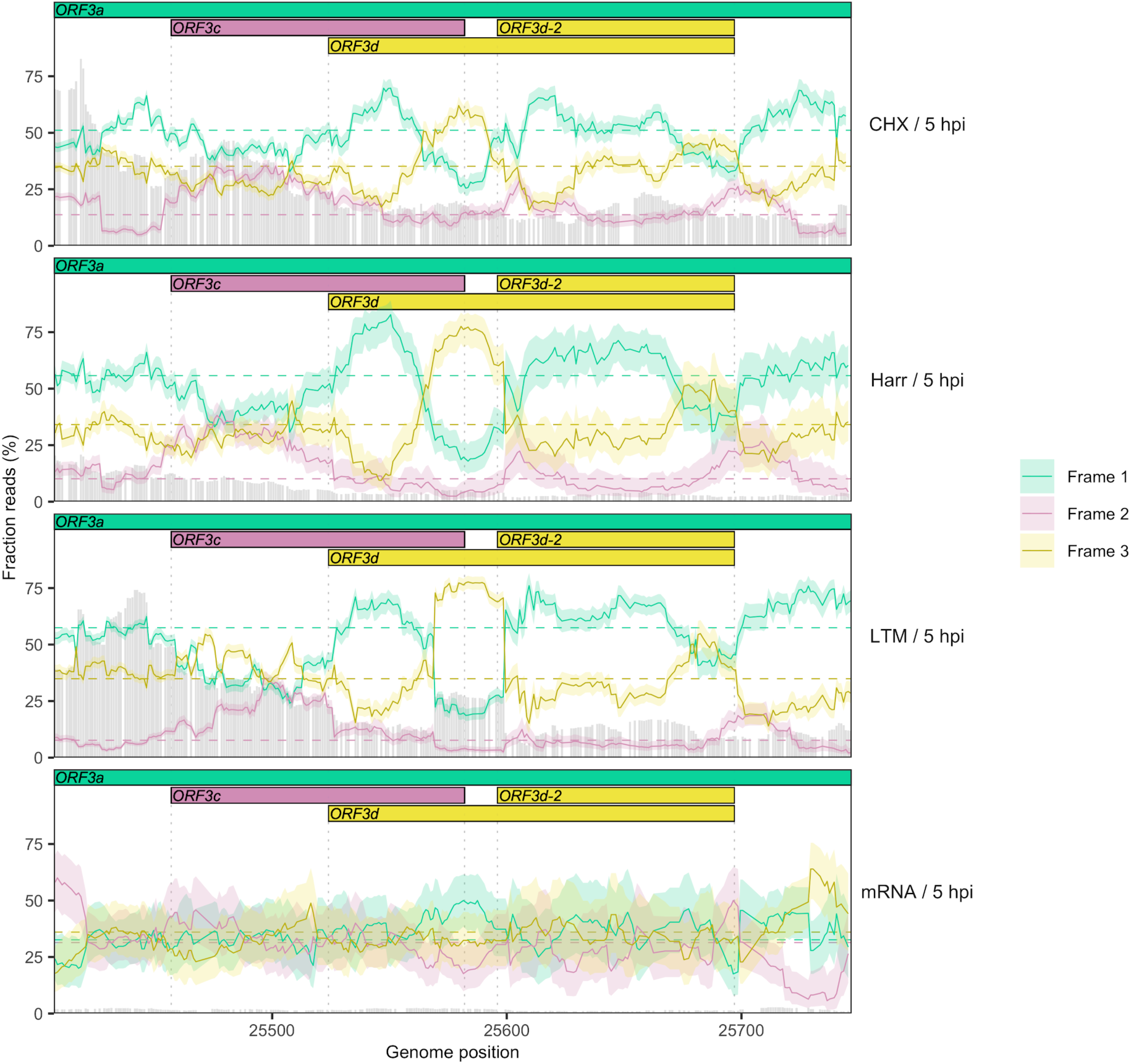
Reading frame of ribosome profiling reads in the *ORF3c*/*ORF3d* region of *ORF3a* for individual treatments. Details as in Figure 2C, except results are shown separately after pooling the two biological replicates for each treatment (CHX=cycloheximide, Harr=harringtonine, LTM=lactimidomycin, mRNA=RNA-seq) (Finkel et al. 2020). Genes and their reading frames are denoted by color according to Figure 1, with Frames 1, 2, and 3 corresponding to the frames beginning at codon positions 1, 2, and 3 of *ORF3a*, respectively. Clear evidence for translation in the appropriate reading frame is observed for *ORF3c* (burgundy) and *ORF3d*/*ORF3d-2* (gold). The status of the first half of *ORF3d* is obscured by the signal of *ORF3c*, but the increase in Frame 1 (*ORF3a*) reads and drop in proportion of Frame 2 (*ORF3c*) reads at the start of *ORF3d* is suggestive of an otherwise-unexpected biological change at this locus.

**Figure 2—figure supplement 6.**
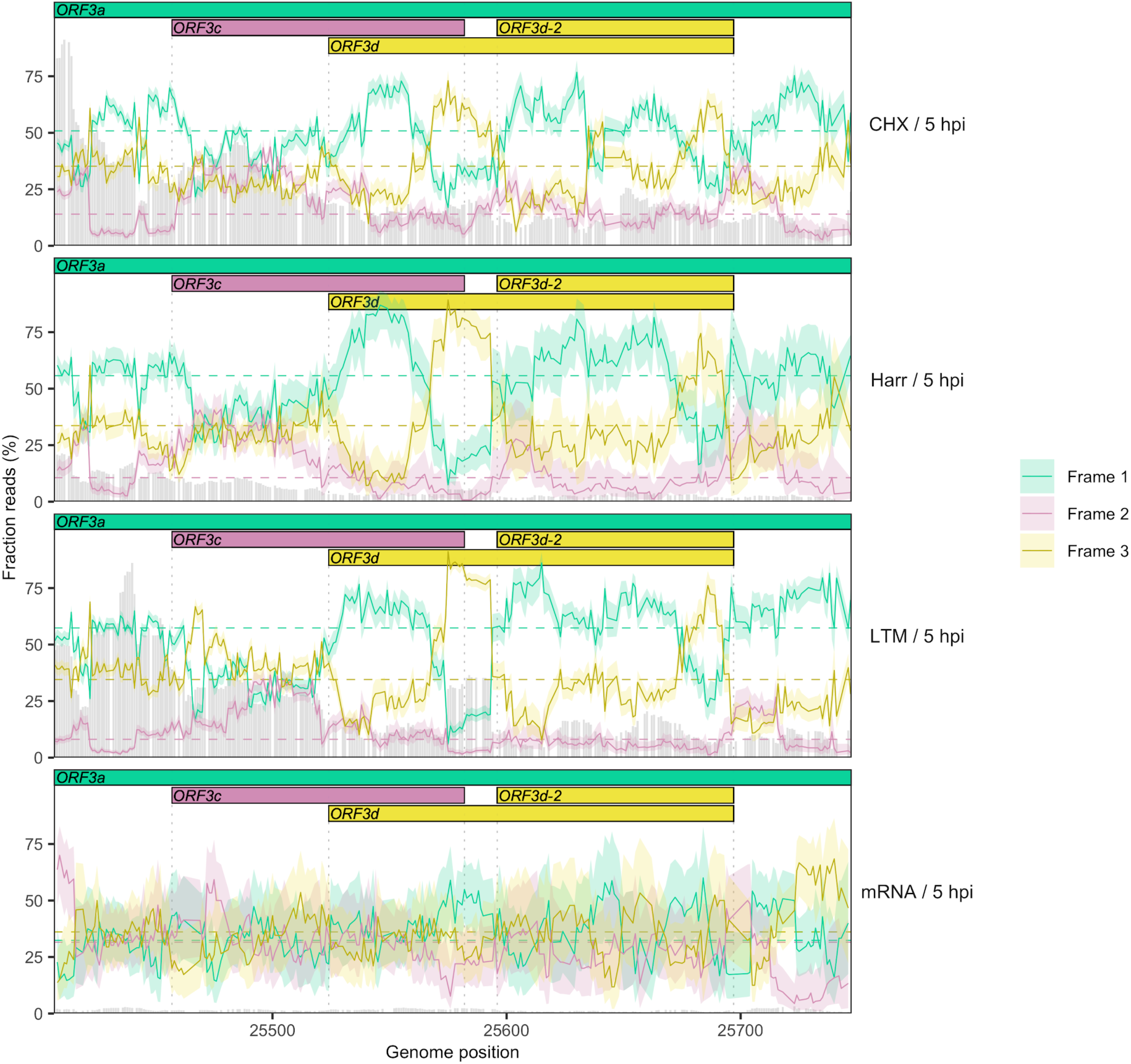
Reading frame of ribosome profiling reads in the *ORF3c*/*ORF3d* region of *ORF3a* for individual treatments using a smaller sliding window. Details as in Figure 2C and Figure 2—figure supplement 5, except results are shown for a sliding window size of 19 nt (step size=1 nt).

**Figure 2—figure supplement 7.**
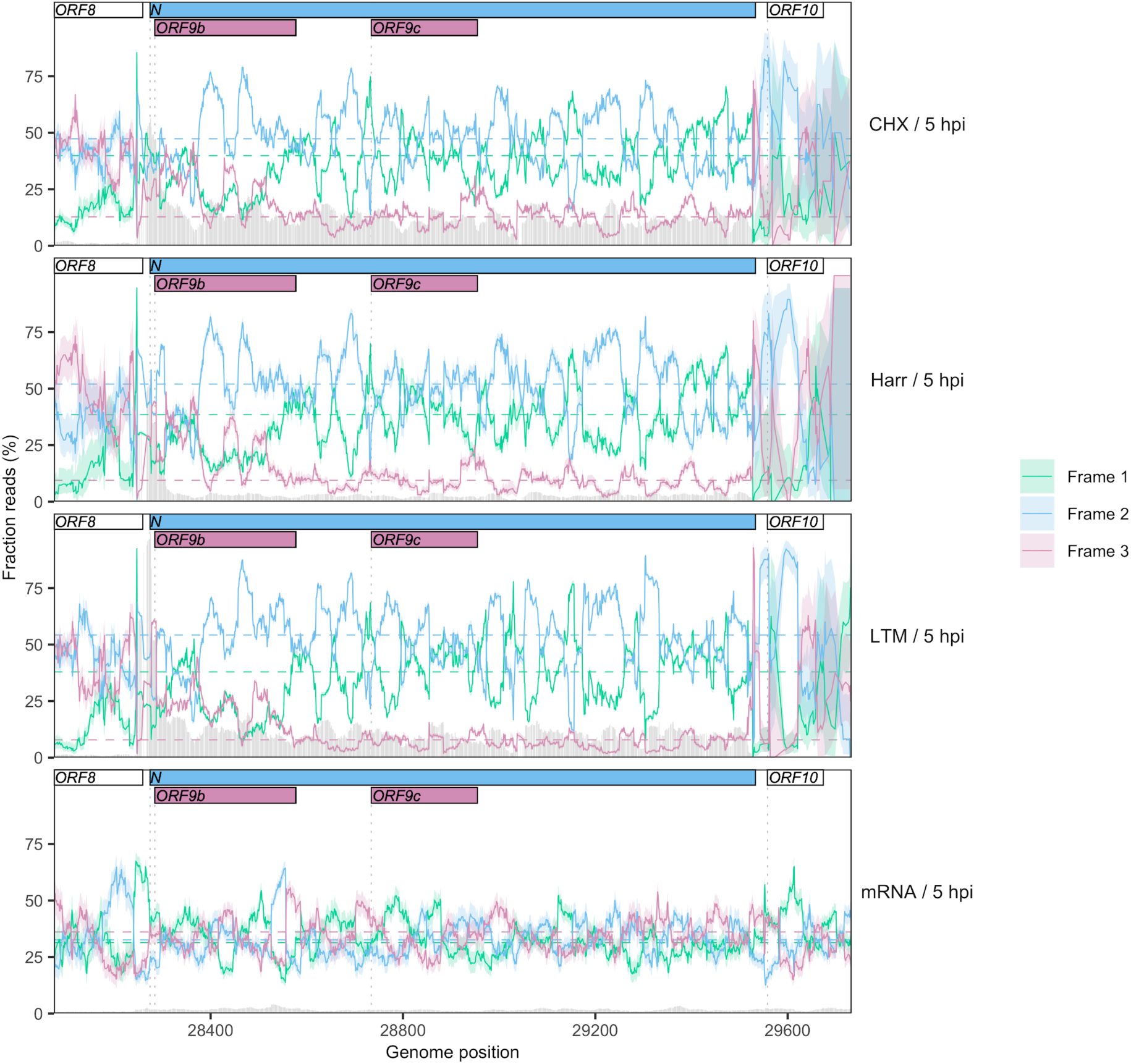
Reading frame of ribosome profiling reads in the *N*/*ORF9b*/*ORF9c* region. Details as in Figure 2C and Figure 2—figure supplement 5, except Frames 1, 2, and 3 correspond to the frames beginning at positions 1, 2, and 3 of the Wuhan-Hu-1 reference genome, respectively. Expression of *ORF9b* is evidenced by a peak in the fraction of reads mapping to Frame 3 (burgundy) at the appropriate region, as well as a peak in read depth (gray bars) immediately upstream (Figure 2A). Translational status of *ORF9c* is ambiguous, with minimal enrichment of reads in the appropriate frame (burgundy) along the gene.

**Figure 2—figure supplement 8.**
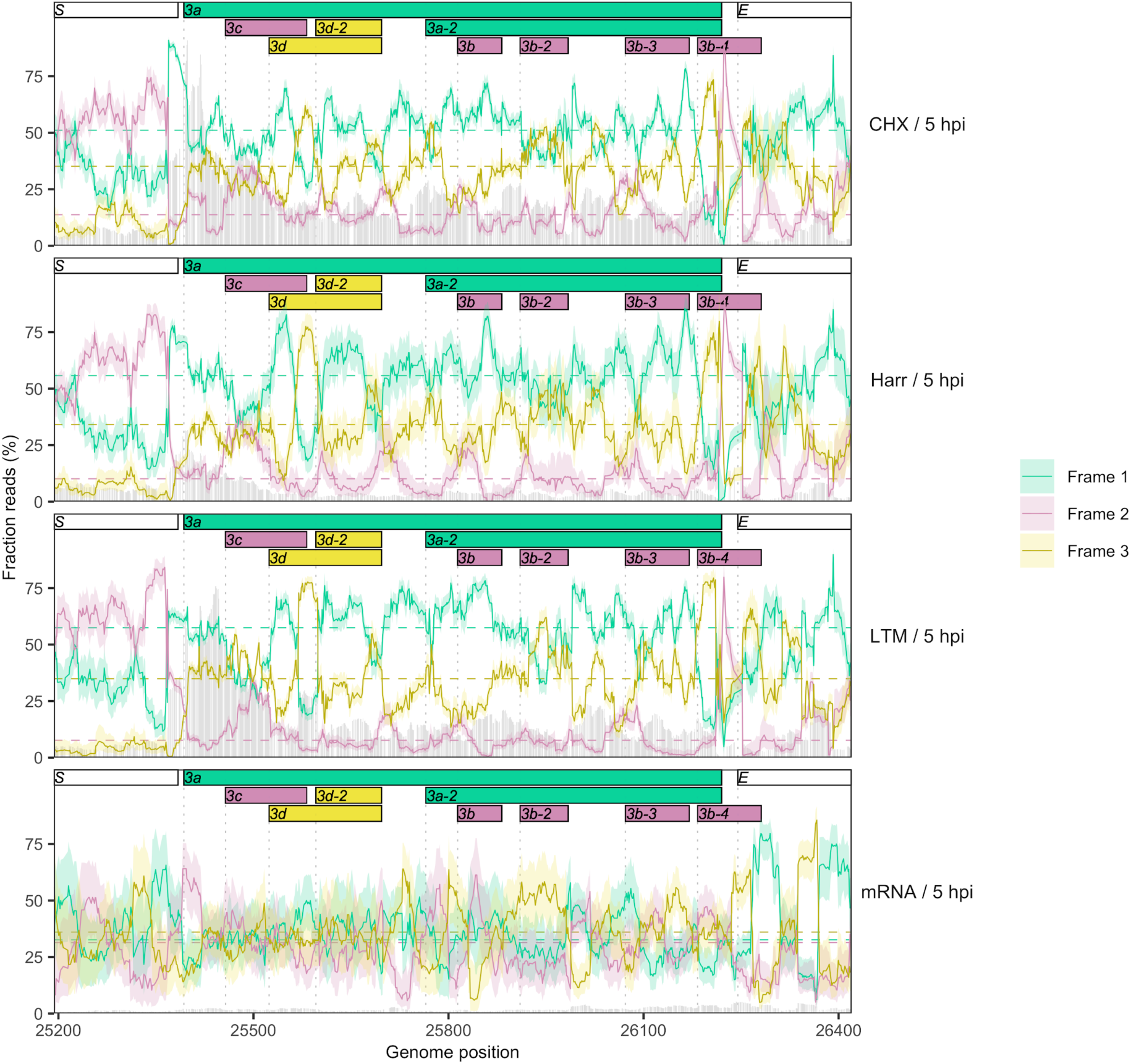
Reading frame of ribosome profiling reads in the full *ORF3a* gene. Details as in Figure 2C and Figure 2—figure supplement 5. No clear evidence is observed for the expression of any of the *ORF3b* ORFs present in SARS-CoV-2, but the translational status of these ORFs requires further investigation.

**Figure 2—figure supplement 9.**
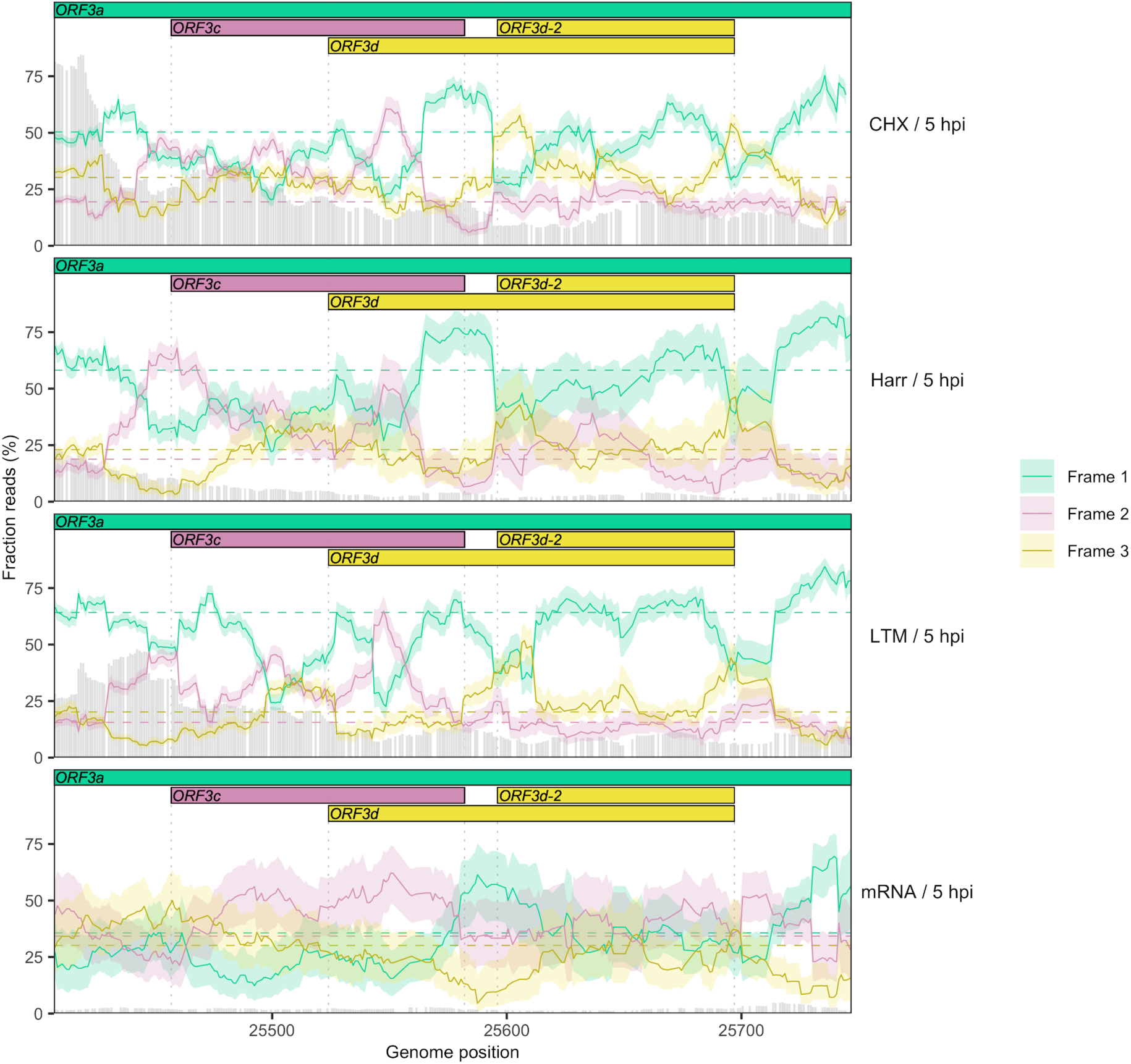
Reading frame of ribosome profiling reads in the *ORF3c*/*ORF3d* region of *ORF3a* for reads of length 29 nt. Details as in Figure 2C and Figure 2—figure supplement 5, except results are shown for reads of length 29 nt, which are also indicative of frame but have slightly lower coverage (Figure 2—figure supplement 3) and exhibit frame-related periodicity. Evidence for *ORF3c* (Frame 2; burgundy) is more pronounced using this read length, while the peak for *ORF3d* (Frame 3; gold) occurs slightly later, overlapping *ORF3d-2*.

**Figure 3—figure supplement 1.**
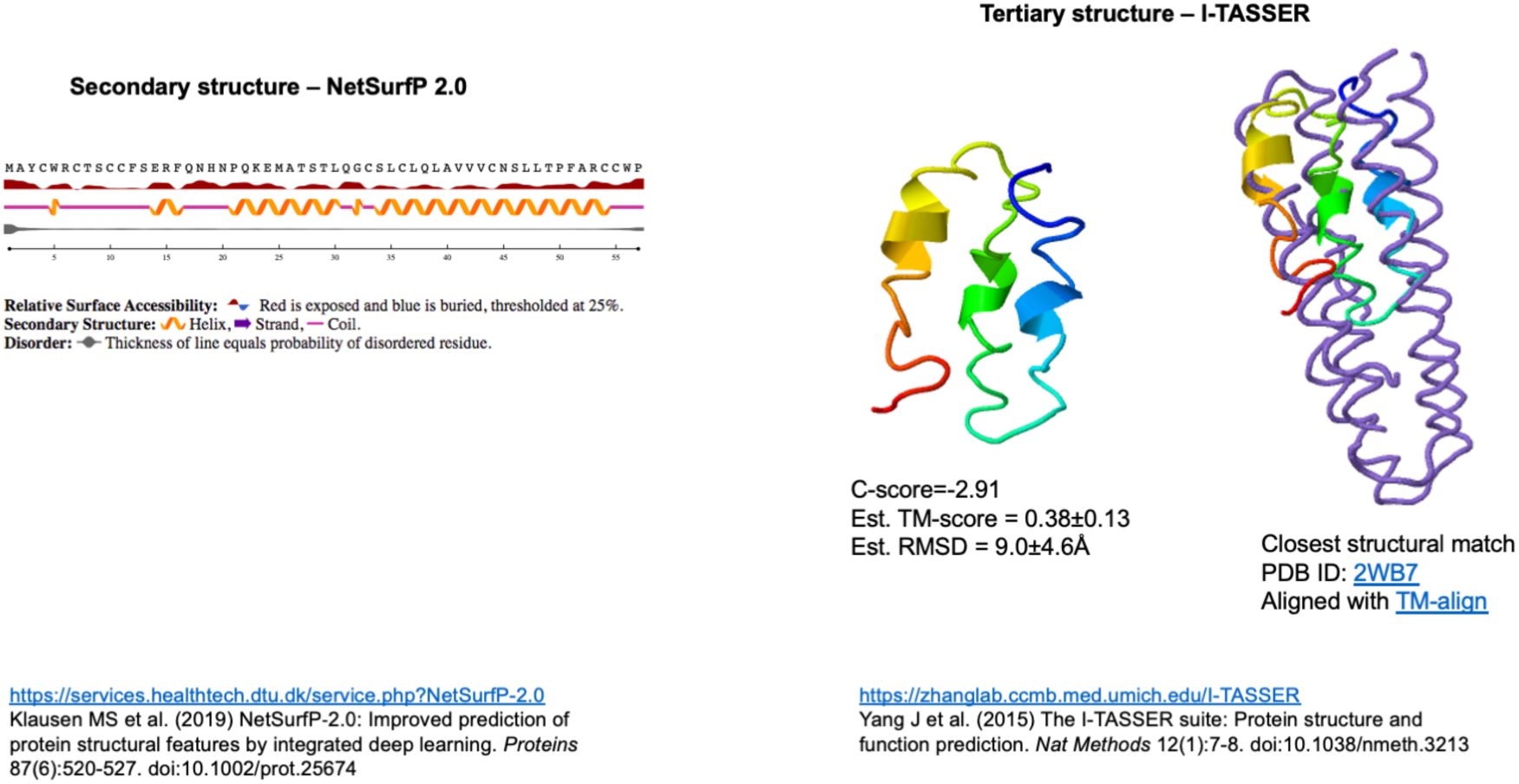
Structural prediction for the ORF3d protein. Independent computational modeling predictions of *α*-helices at the secondary (left inset, carried out in NetSurf v2) and tertiary structure levels (right inset, carried out in I-TASSER). Folding concordance with the closest protein structure match is shown (rightmost inset, aligned with TM-align). For explanation of shown metrics, see https://zhanglab.ccmb.med.umich.edu/I-TASSER. Chan et al. (2020) also predict a fold with α-helices (Raven Kok, pers. comm.).

**Figure 3—figure supplement 2.**
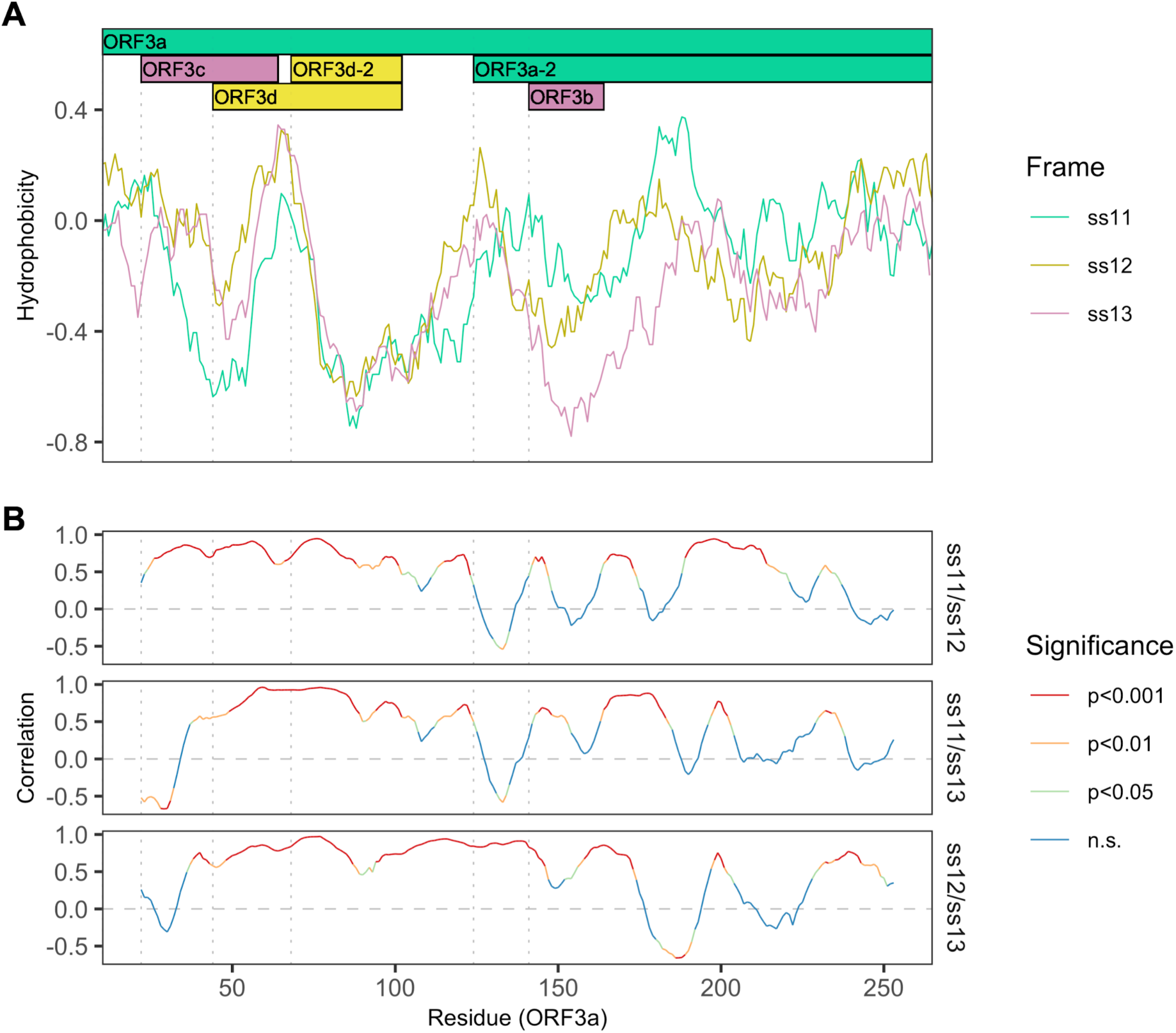
Correlations between hydrophobicity profiles of amino acid sequences encoded by the three forward-strand reading frames of the *ORF3a* region. Vertical dotted lines denote gene start sites. **(A)** Hydrophobicity profiles of the amino acid sequences encoded by all three forward-strand reading frames of *ORF3a*. Details as in Figure 3B, except the entire region encoding *ORF3a* is shown. **(B)** Correlations (Spearman’s rank) between hydrophobicity profiles for the three forward-strand reading frames. Color denotes significance.

**Figure 3—figure supplement 3.**
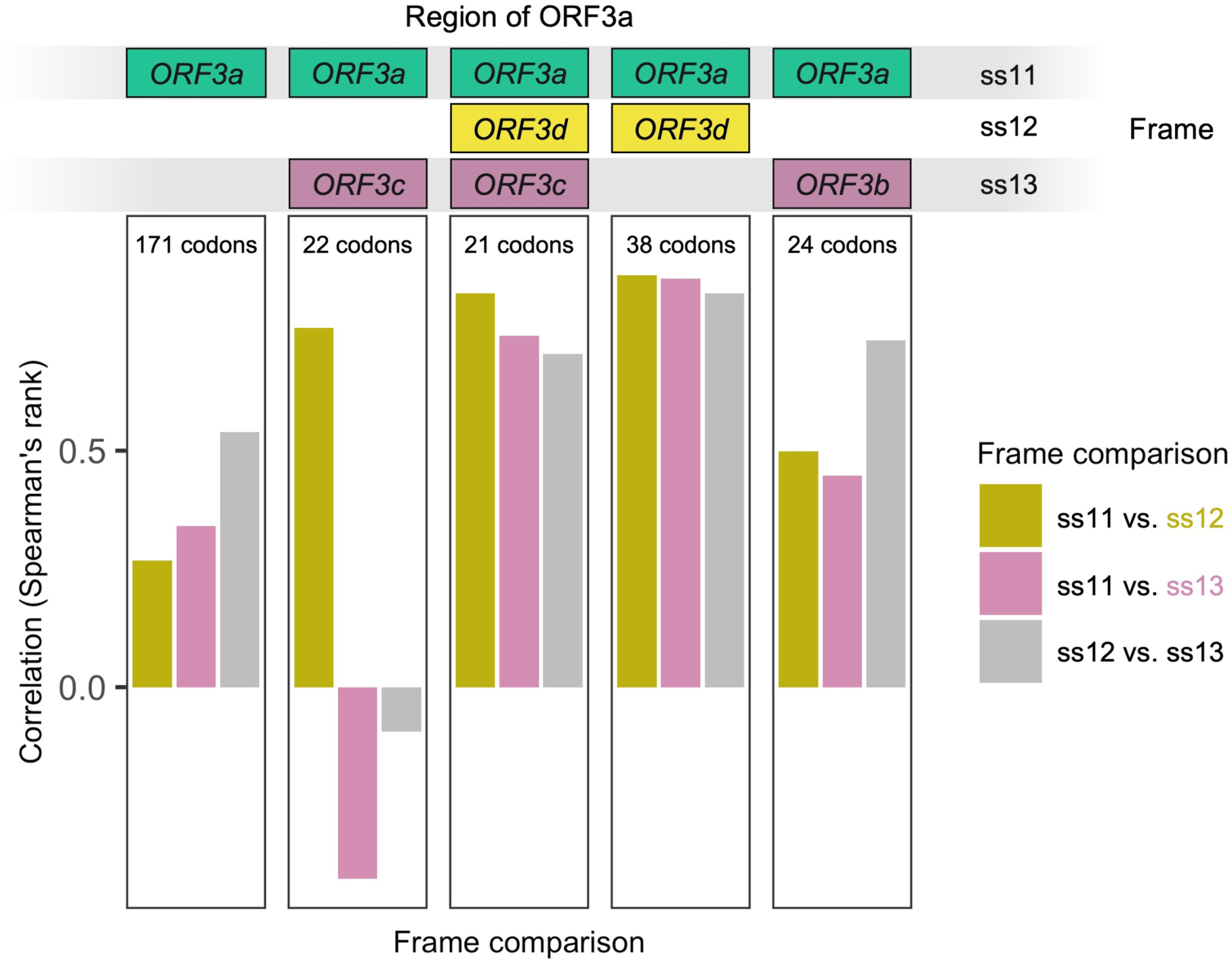
Correlation between hydrophobicity profiles in the amino acid sequences encoded by all three forward-strand frames of *ORF3a* by gene subregion. Details as in Figure 3—figure supplement 2. *ORF3a* (left) refers to those codons of *ORF3a* that encode residues not overlapping any hypothesized overlapping genes. Codons counts refer to *ORF3a*, i.e, 21 codons of *ORF3a* (20 codons of *ORF3d*) are involved in a triple overlap (at least one nucleotide) with *ORF3c*.

**Figure 5—figure supplement 1.**
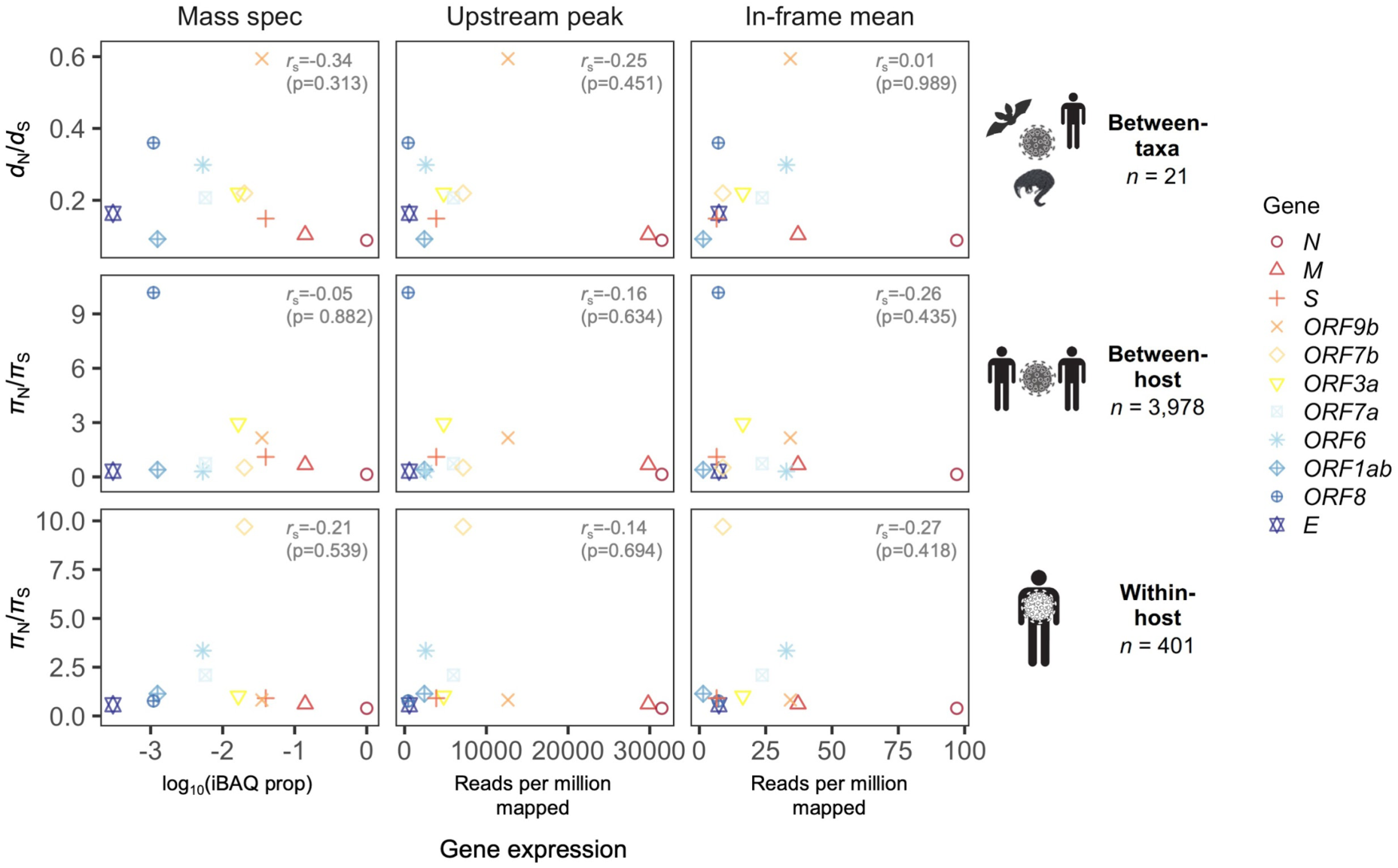
Correlation between natural selection and gene expression. A weak negative Spearman correlation between *d*_N_/*d*_S_ and gene expression is observed across all three evolutionary levels (between-taxa, between-host, and within-host). The x axes of different panels differ by method for inferring gene expression, which are those shown in Figure 2B: (1) ‘Mass spec’ refers to log_10_(iBAQ prop), where iBAQ prop is the proportion of the maximum intensity-based absolute quantification (iBAQ) observed; (2) ‘Upstream peak’ is the peak in ribosome profiling read depth upstream of the gene start site, measured for reads of length 29-31nt (Figure 2A); and (3) ‘In-frame mean’ is the mean read depth at Codon pos 1, measured for reads of length 30nt (Figure 2B). A given gene (color) has only one selection value (y-axis) per evolutionary level, and only one gene expression value (x-axis) per method for determining expression. Selection was calculated as *d*_N_/*d*_S_ (*π*_N_/*π*_S_) except for the overlapping gene *ORF9b*, where the OLG-appropriate measure *d*_NN_/*d*_NS_ (*π*_NN_/*π*_NS_) was used (see Methods).

**Figure 5—figure supplement 2.**
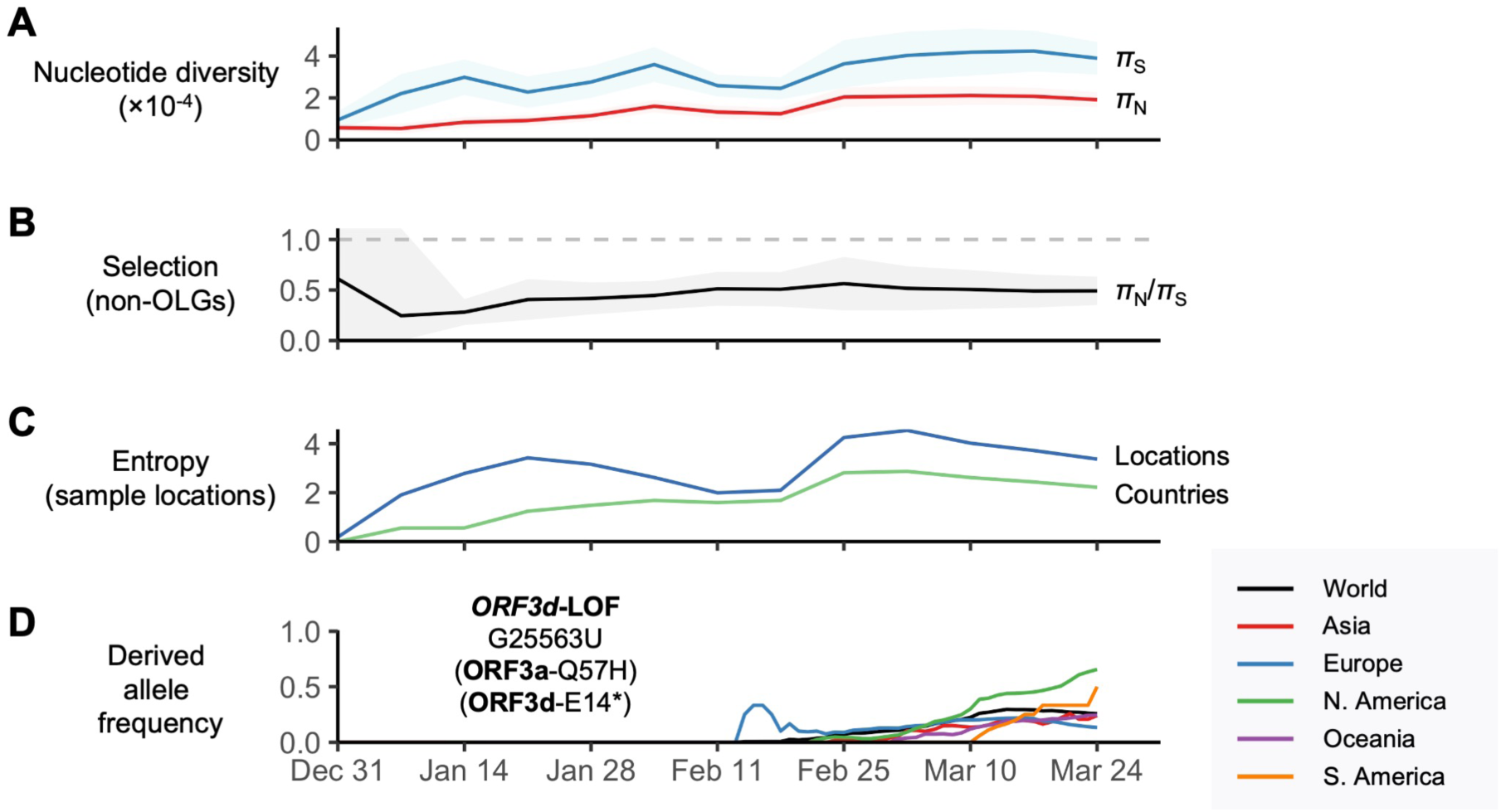
SARS-CoV-2 between-host nucleotide diversity and allele frequencies as a function of time. Analyses refer to human SARS-CoV-2 consensus sequences obtained from GISAID. Results show sliding windows of 14 days (step size=7 days) representing 13 time points since the first GISAID sample was collected (EPI_ISL_402123 on 12/24/2019). Regions with overlapping genes were excluded from estimates of *π*. Shaded regions show standard error of mean *π*_N_, *π*_S_, or *π*_N_/*π*_S_ (10,000 bootstrap replicates, codon unit). **(A)** Nonsynonymous (*π*_N_) and synonymous (*π*_S_) nucleotide diversity. **(B)** *π*_N_/*π*_S_, where the horizontal dashed gray line denotes the *π*_N_/*π*_S_ ratio expected under neutrality (1.0). **(C)** Diversity of sampling locations measured as entropy, defined as -∑*p*\*ln(*p*), where *p* is the number of distinct (unique) locations or countries reported for a given window (Ewens and Grant 2001). **(D)** Allele frequency trajectories of the *ORFd*-LOF mutation, colored by continent and limited to sufficient sample sizes (Supplementary Table 19).

**Figure 6—figure supplement 1.**
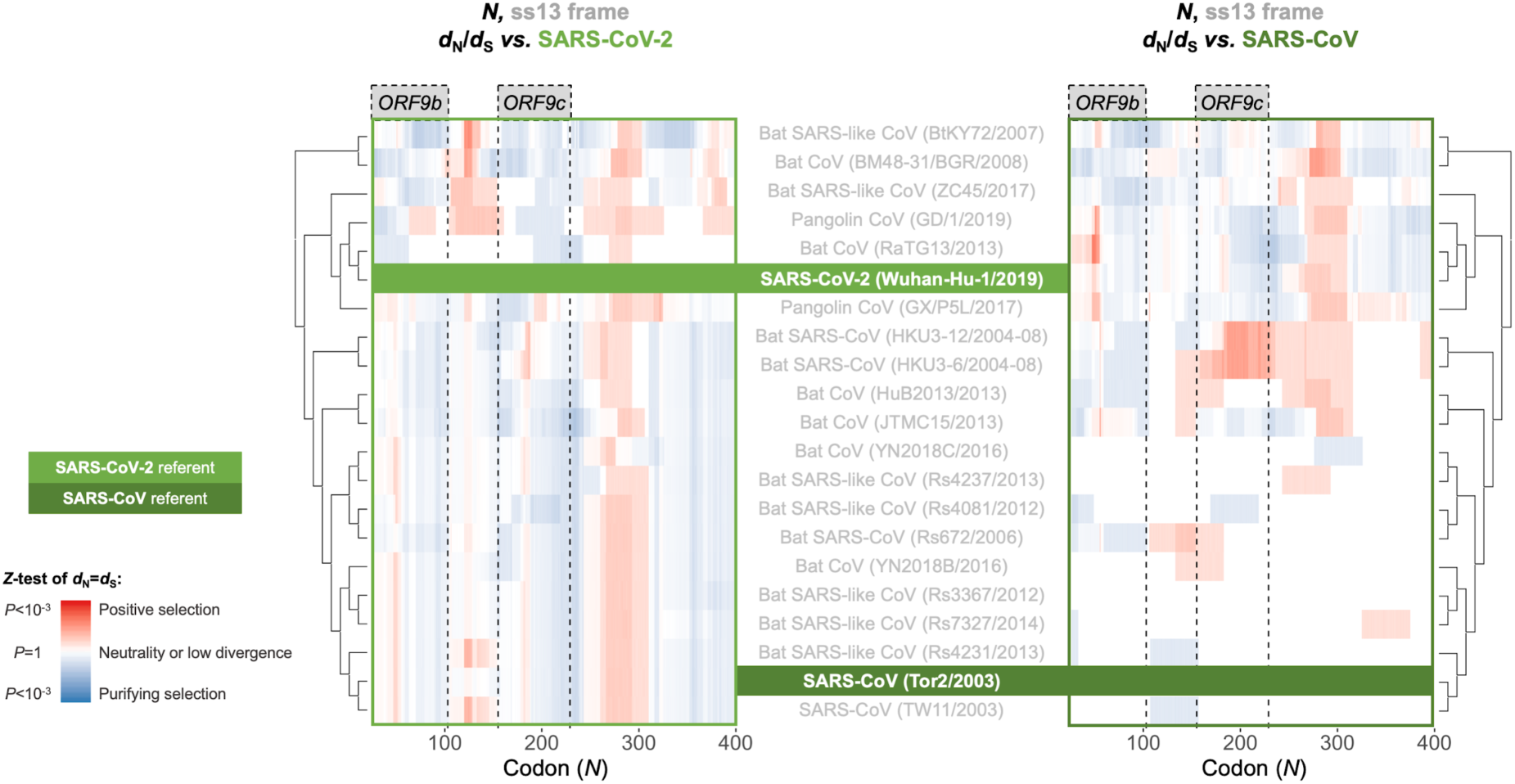
Between-taxa sliding window of genes overlapping *N*. Pairwise OLGenie analysis of the N gene’s ss13 reading frame across members of the species *Severe acute respiratory syndrome-related coronavirus*. Details as in Figure 6, except only the ss13 frame is shown.

**Figure 8—figure supplement 1.**
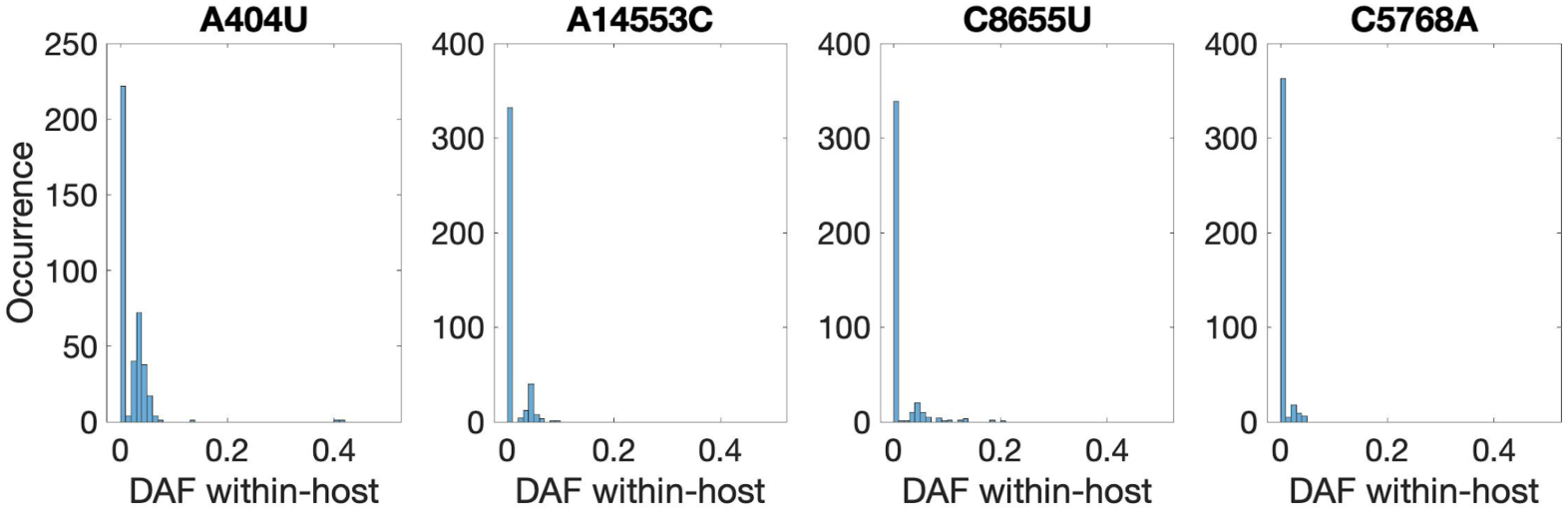
Recurrent nonsynonymous mutations observed in multiple human hosts. Histogram of the derived allele frequency (DAF) of the four most common recurrent within-host protein-coding mutations across 401 samples (bin width=0.01). Mutation A404U introduces a premature stop codon in *nsp1*, whereas the remainder are nonsynonymous.

## Notes

https://github.com/chasewnelson/SARS-CoV-2-ORF3d

https://zenodo.org/record/4052729

### Competing Interest Statement

The authors have declared no competing interest.

### Summary of Updates

Updated nomenclature to reflect community consensus, specifically by naming the novel overlapping gene ORF3d instead of ORF3c, and using the species name Severe acute respiratory syndrome-related coronavirus instead of the more vague, general, subgenus-reminiscent term 'sarbecovirus'. Addressed reviewer comments by using the reviewer-suggested title; thoroughly updating the ribosome profiling analysis; including OLGs ORF3c and ORF3b in analyses; adding a hydrophobicity profile analysis; improving figures for clarity; and general edits. Added links to GitHub and Zenodo for supplementary material.

## References

Affram Y, Zapata JC, Gholizadeh Z, Tolbert WD, Zhou W, Iglesias-Ussel MD, Pazgier M, Ray K, Latinovic OS, Romerio F. 2019. The HIV-1 Antisense Protein ASP Is a Transmembrane Protein of the Cell Surface and an Integral Protein of the Viral Envelope. J Virol 93:e00574–19.

Atchley WR, Zhao J, Fernandes AD, Druke T. 2005. Solving the protein sequence metric problem. Proceedings of the National Academy of Sciences 102:6395–6400.

Bartonek L, Braun D, Zagrovic B. 2020. Frameshifting preserves key physicochemical properties of proteins. Proc Natl Acad Sci USA 117:5907–5912.

Bartonek L, Zagrovic B. 2019. VOLPES: an interactive web-based tool for visualizing and comparing physicochemical properties of biological sequences. Nucleic Acids Research 47:W632–W635.

Benjamini Y, Hochberg Y. 1995. Controlling the false discovery rate: a practical and powerful approach to multiple testing. Journal of the Royal Statistical Society B 57:289–300.

Benjamini Y, Yekutieli D. 2001. The control of the false discovery rate in multiple testing under dependency. The Annals of Statistics 29:1165–1188.

Bezstarosti K, Lamers MM, Haagmans BL, Demmers JAA. 2020. Targeted Proteomics for the Detection of SARS-CoV-2 Proteins. Biochemistry Available from: http://biorxiv.org/lookup/doi/10.1101/2020.04.23.057810

Bojkova D, Klann K, Koch B, Widera M, Krause D, Ciesek S, Cinatl J, Münch C. 2020. SARS-CoV-2 infected host cell proteomics reveal potential therapy targets. In Review Available from: https://www.researchsquare.com/article/rs-17218/v1

Boni MF, Lemey P, Jiang X, Lam TT-Y, Perry B, Castoe T, Rambaut A, Robertson DL. 2020. Evolutionary origins of the SARS-CoV-2 sarbecovirus lineage responsible for the COVID-19 pandemic. Nature Microbiology, doi: 10.1038/s41564-020-0771-4

Boussau B, Gouy M. 2006. Efficient likelihood computations with nonreversible models of evolution. Syst Biol 55:756–768.

Cagliani R, Forni D, Clerici M, Sironi M. 2020. Coding potential and sequence conservation of SARS-CoV-2 and related animal viruses. Infection, Genetics and Evolution 83:104353.

Calviello L, Ohler U. 2017. Beyond Read-Counts: Ribo-seq Data Analysis to Understand the Functions of the Transcriptome. Trends in Genetics 33:728–744.

Cassan E, Arigon-Chifolleau A-M, Mesnard J-M, Gross A, Gascuel O. 2016. Concomitant emergence of the antisense protein gene of HIV-1 and of the pandemic. Proc Natl Acad Sci USA 113:11537–11542.

Chan JF-W, Kok K-H, Zhu Z, Chu H, To KK-W, Yuan S, Yuen K-Y. 2020. Genomic characterization of the 2019 novel human-pathogenic coronavirus isolated from a patient with atypical pneumonia after visiting Wuhan. Emerging Microbes & Infections 9:221–236.

Cock PJA, Antao T, Chang JT, Chapman BA, Cox CJ, Dalke A, Friedberg I, Hamelryck T, Kauff F, Wilczynski B, et al. 2009. Biopython: freely available Python tools for computational molecular biology and bioinformatics. Bioinformatics 25:1422–1423.

Cui J, Li F, Shi Z-L. 2019. Origin and evolution of pathogenic coronaviruses. Nature Reviews Microbiology 17:181–192.

Daugherty MD, Malik HS. 2012. Rules of Engagement: Molecular Insights from Host-Virus Arms Races. Annual Review of Genetics 46:677–700.

Davidson AD, Williamson MK, Lewis S, Shoemark D, Carroll MW, Heesom KJ, Zambon M, Ellis J, Lewis PA, Hiscox JA, et al. 2020. Characterisation of the transcriptome and proteome of SARS-CoV-2 reveals a cell passage induced in-frame deletion of the furin-like cleavage site from the spike glycoprotein. Genome Med 12:68.

Degner JF, Marioni JC, Pai AA, Pickrell JK, Nkadori E, Gilad Y, Pritchard JK. 2009. Effect of read-mapping biases on detecting allele-specific expression from RNA-sequencing data. Bioinformatics 25:3207–3212.

Do CB, Mahabhashyam MSP, Brudno M, Batzoglou S. 2005. ProbCons: Probabilistic consistency-based multiple sequence alignment. Genome Research 15:330–340.

Ewens WJ, Grant GR. 2001. Statistical Methods in Bioinformatics. New York: Springer-Verlag

Fernandes JD, Faust TB, Strauli NB, Smith C, Crosby DC, Nakamura RL, Hernandez RD, Frankel AD. 2016. Functional Segregation of Overlapping Genes in HIV. Cell 167:1762–1773.

Finkel Y, Mizrahi O, Nachshon A, Weingarten-Gabbay S, Morgenstern D, Yahalom-Ronen Y, Tamir H, Achdout H, Stein D, Israeli O, et al. 2020. The coding capacity of SARS-CoV-2. Nature, in press, https://doi.org/10.1038/s41586-020-2739-1

Firth AE. 2020. A putative new SARS-CoV protein, 3c, encoded in an ORF overlapping ORF3a. Journal of General Virology, https://doi.org/10.1099/jgv.0.001469

Flynn JA, Purushotham D, Choudhary MNK, Zhou X, Fan C, Matt G, Li D, Wang T. 2020. Exploring the coronavirus pandemic with the WashU Virus Genome Browser. Nat Genet [Internet]. Available from: http://www.nature.com/articles/s41588-020-0697-z

Forni D, Cagliani R, Clerici M, Sironi M. 2017. Molecular Evolution of Human Coronavirus Genomes. Trends in Microbiology 25:35–48.

Forster P, Forster L, Renfrew C, Forster M. 2020. Phylogenetic network analysis of SARS-CoV-2 genomes. Proc Natl Acad Sci USA:202004999.

Fung S-Y, Yuen K-S, Ye Z-W, Chan C-P, Jin D-Y. 2020. A tug-of-war between severe acute respiratory syndrome coronavirus 2 and host antiviral defence: lessons from other pathogenic viruses. Emerging Microbes & Infections 9:558–570.

Gazave E, Chang D, Clark AG, Keinan A. 2013. Population Growth Inflates the Per-Individual Number of Deleterious Mutations and Reduces Their Mean Effect. Genetics 195:969–978.

Ge H, Wang X, Yuan X, Xiao G, Wang C, Deng T, Yuan Q, Xiao X. 2020. The epidemiology and clinical information about COVID-19. European Journal of Clinical Microbiology & Infectious Diseases 39:1011–1019.

Gorbalenya AE, Enjuanes L, Ziebuhr J, Snijder EJ. 2006. Nidovirales: Evolving the largest RNA virus genome. Virus Research 117:17–37.

Gorbalenya AE, Baker SC, Baric RS, de Groot RJ, Drosten C, Gulyaeva AA, Haagmans BL, Lauber C, Leontovich AM, Neuman BW, et al. 2020. The species Severe acute respiratory syndrome-related coronavirus: classifying 2019-nCoV and naming it SARS-CoV-2. Nature Microbiology 5:536–544.

Gordon DE, Jang GM, Bouhaddou M, Xu J, Obernier K, White KM, O’Meara MJ, Rezelj VV, Guo JZ, Swaney DL, et al. 2020. A SARS-CoV-2 protein interaction map reveals targets for drug repurposing. Nature [Internet]. Available from: http://www.nature.com/articles/s41586-020-2286-9

Greenbaum J, Sidney J, Chung J, Brander C, Peters B, Sette A. 2011. Functional classification of class II human leukocyte antigen (HLA) molecules reveals seven different supertypes and a surprising degree of repertoire sharing across supertypes. Immunogenetics 63:325–335.

Grifoni A, Sidney J, Zhang Y, Scheuermann RH, Peters B, Sette A. 2020. A Sequence Homology and Bioinformatic Approach Can Predict Candidate Targets for Immune Responses to SARS-CoV-2. Cell Host & Microbe 27:671–680.e2.

Grubaugh ND, Gangavarapu K, Quick J, Matteson NL, De Jesus JG, Main BJ, Tan AL, Paul LM, Brackney DE, Grewal S, et al. 2019. An amplicon-based sequencing framework for accurately measuring intrahost virus diversity using PrimalSeq and iVar. Genome Biol 20:8.

Hachim A, Kavian N, Cohen CA, Chin AWH, Chu DKW, Mok CKP, Tsang OTY, Yeung YC, Perera RAPM, Poon LLM, et al. 2020. ORF8 and ORF3b antibodies are accurate serological markers of early and late SARS-CoV-2 infection. Nat Immunol 21:1293– 1301.

Helmy YA, Fawzy M, Elaswad A, Sobieh A, Kenney SP, Shehata AA. 2020. The COVID-19 Pandemic: A Comprehensive Review of Taxonomy, Genetics, Epidemiology, Diagnosis, Treatment, and Control. JCM 9:1225.

Holmes EC. 2009. The Evolution and Emergence of RNA Viruses. New York: Oxford University Press.

Holmes EC, Lipman DJ, Zamarin D, Yewdell JW. 2006. Comment on “Large-Scale Sequence Analysis of Avian Influenza Isolates.” Science 313:1573b–1573b.

Hughes AL, Friedman R, Glenn NL. 2006. The Future of Data Analysis in Evolutionary Genomics. Current Genomics 7:227–234.

Ingolia NT, Ghaemmaghami S, Newman JRS, Weissman JS. 2009. Genome-Wide Analysis in Vivo of Translation with Nucleotide Resolution Using Ribosome Profiling. Science 324:218–223.

Jukes TH, Cantor CR. 1969. Evolution of Protein Molecules. In: Munro HN, editor. Mammalian Protein Metabolism. New York: Academic Press. p. 21–132. Available from: https://linkinghub.elsevier.com/retrieve/pii/B9781483232119500097

Jungreis I, Sealfon R, Kellis M. 2020. Sarbecovirus comparative genomics elucidates gene content of SARS-CoV-2 and functional impact of COVID-19 pandemic mutations. Genomics Available from: http://biorxiv.org/lookup/doi/10.1101/2020.06.02.130955

Jurtz V, Paul S, Andreatta M, Marcatili P, Peters B, Nielsen M. 2017. NetMHCpan-4.0: Improved Peptide–MHC Class I Interaction Predictions Integrating Eluted Ligand and Peptide Binding Affinity Data. J.I. 199:3360–3368.

Katoh K, Standley DM. 2013. MAFFT Multiple Sequence Alignment Software Version 7: Improvements in Performance and Usability. Molecular Biology and Evolution 30:772–780.

Keese PK, Gibbs A. 1992. Origins of genes: “big bang” or continuous creation? Proceedings of the National Academy of Sciences 89:9489–9493.

Konno Y, Kimura I, Uriu K, Fukushi M, Irie T, Koyanagi Y, Sauter D, Gifford RJ, Nakagawa S, Sato K. 2020. SARS-CoV-2 ORF3b Is a Potent Interferon Antagonist Whose Activity Is Increased by a Naturally Occurring Elongation Variant. Cell Reports 32:108185.

Kopecky-Bromberg SA, Martínez-Sobrido L, Frieman M, Baric RA, Palese P. 2007. Severe Acute Respiratory Syndrome Coronavirus Open Reading Frame (ORF) 3b, ORF 6, and Nucleocapsid Proteins Function as Interferon Antagonists. JVI 81:548–557.

Korber B, Fischer WM, Gnanakaran S, Yoon H, Theiler J, Abfalterer W, Hengartner N, Giorgi EE, Bhattacharya T, Foley B, et al. 2020. Tracking Changes in SARS-CoV-2 Spike: Evidence that D614G Increases Infectivity of the COVID-19 Virus. Cell 182:812–827.e19.

Kosakovsky-Pond SL. 2020. Natural selection analysis of SARS-CoV-2/COVID-19. usegalaxy [Internet]. Available from: https://covid19.galaxyproject.org/evolution/

Lam TT-Y, Shum MH-H, Zhu H-C, Tong Y-G, Ni X-B, Liao Y-S, Wei W, Cheung WY-M, Li W-J, Li L-F, et al. 2020. Identifying SARS-CoV-2 related coronaviruses in Malayan pangolins. Nature [Internet] in press. Available from: http://www.nature.com/articles/s41586-020-2169-0

Langmead B, Salzberg SL. 2012. Fast gapped-read alignment with Bowtie 2. Nat Methods 9:357–359.

Langmead B, Wilks C, Antonescu V, Charles R. 2019. Scaling read aligners to hundreds of threads on general-purpose processors.Hancock J, editor. Bioinformatics 35:421– 432.

Larsson A. 2014. AliView: a fast and lightweight alignment viewer and editor for large datasets. Bioinformatics 30:3276–3278.

Lawrence M, Huber W, Pagès H, Aboyoun P, Carlson M, Gentleman R, Morgan MT, Carey VJ. 2013. Software for Computing and Annotating Genomic Ranges.Prlic A, editor. PLoS Comput Biol 9:e1003118.

Lokugamage KG, Hage A, Schindewolf C, Rajsbaum R, Menachery VD. 2020. SARS-CoV-2 is sensitive to type I interferon pretreatment. Microbiology Available from: http://biorxiv.org/lookup/doi/10.1101/2020.03.07.982264

McBride R, Fielding B. 2012. The Role of Severe Acute Respiratory Syndrome (SARS)- Coronavirus Accessory Proteins in Virus Pathogenesis. Viruses 4:2902–2923.

McKinney W. 2010. Data structures for statistical computing in python. In: Proceedings of the 9th Python in Science Conference. Vol. 445. p. 51–56.

Michel CJ, Mayer C, Poch O, Thompson JD. 2020. Characterization of accessory genes in coronavirus genomes. Virol J 17:131.

Nei M, Gojobori T. 1986. Simple methods for estimating the numbers of synonymous and nonsynonymous nucleotide substitutions. Molecular Biology and Evolution 3:418– 426.

Nei M, Kumar S. 2000. Molecular Evolution and Phylogenetics. New York, NY: Oxford University Press

Nelson CW, Ardern Z, Wei X. 2020. OLGenie: Estimating Natural Selection to Predict Functional Overlapping Genes. Molecular Biology and Evolution in press:msaa087.

Nelson CW, Hughes AL. 2015. Within-host nucleotide diversity of virus populations: Insights from next-generation sequencing. Infection, Genetics and Evolution 30:1–7.

Nelson CW, Moncla LH, Hughes AL. 2015. SNPGenie: estimating evolutionary parameters to detect natural selection using pooled next-generation sequencing data. Bioinformatics 31:3709–3711.

Nguyen L-T, Schmidt HA, von Haeseler A, Minh BQ. 2015. IQ-TREE: A Fast and Effective Stochastic Algorithm for Estimating Maximum-Likelihood Phylogenies. Molecular Biology and Evolution 32:268–274.

Paul S, Sidney J, Sette A, Peters B. 2016. TepiTool: A Pipeline for Computational Prediction of T Cell Epitope Candidates. Current Protocols in Immunology 114:18.19.1-18.19.24.

R Core Team. 2018. R: A language and environment for statistical computing. Vienna, Austria: R Foundation for Statistical Computing Available from: https://www.R-project.org/

Rehman S ur, Shafique L, Ihsan A, Liu Q. 2020. Evolutionary Trajectory for the Emergence of Novel Coronavirus SARS-CoV-2. Pathogens 9:240.

Reynisson B, Barra C, Kaabinejadian S, Hildebrand WH, Peters B, Nielsen M. 2020. Improved Prediction of MHC II Antigen Presentation through Integration and Motif Deconvolution of Mass Spectrometry MHC Eluted Ligand Data. Journal of Proteome Research 19:2304–2315.

Rothe C, Schunk M, Sothmann P, Bretzel G, Froeschl G, Wallrauch C, Zimmer T, Thiel V, Janke C, Guggemos W, et al. 2020. Transmission of 2019-nCoV Infection from an Asymptomatic Contact in Germany. N Engl J Med 382:970–971.

Sabath N, Landan G, Graur D. 2008. A Method for the Simultaneous Estimation of Selection Intensities in Overlapping Genes. PLoS ONE 3:e3996.

Schirmer M, D’Amore R, Ijaz UZ, Hall N, Quince C. 2016. Illumina error profiles: resolving fine-scale variation in metagenomic sequencing data. BMC Bioinformatics 17:125.

Schlub TE, Buchmann JP, Holmes EC. 2018. A simple method to detect candidate overlapping genes in viruses using single genome sequences. Molecular Biology and Evolution 35:2572–2581.

Schwanhäusser B, Busse D, Li N, Dittmar G, Schuchhardt J, Wolf J, Chen W, Selbach M. 2011. Global quantification of mammalian gene expression control. Nature 473:337– 342.

Sidney J, Peters B, Frahm N, Brander C, Sette A. 2008. HLA class I supertypes: a revised and updated classification. BMC Immunology 9:1.

Soubrier J, Steel M, Lee MSY, Der Sarkissian C, Guindon S, Ho SYW, Cooper A. 2012. The Influence of Rate Heterogeneity among Sites on the Time Dependence of Molecular Rates. Molecular Biology and Evolution 29:3345–3358.

Suyama M, Torrents D, Bork P. 2006. PAL2NAL: robust conversion of protein sequence alignments into the corresponding codon alignments. Nucleic Acids Research 34:W609–W612.

Tavaré S. 1986. Some probabilistic and statistical problems in the analysis of DNA sequences. Lectures on Mathematics in the Life Sciences 17:57–86.

Tyanova S, Temu T, Cox J. 2016. The MaxQuant computational platform for mass spectrometry-based shotgun proteomics. Nat Protoc 11:2301–2319.

Warren AS, Archuleta J, Feng W, Setubal JC. 2010. Missing genes in the annotation of prokaryotic genomes. BMC Bioinformatics 11:131.

Wei X, Zhang J. 2015. A Simple Method for Estimating the Strength of Natural Selection on Overlapping Genes. Genome Biology and Evolution 7:381–390.

Wells HL, Letko M, Lasso G, Ssebide B, Nziza J, Byarugaba DK, Navarrete-Macias I, Liang E, Cranfield M, Han BA, et al. 2020. The evolutionary history of ACE2 usage within the coronavirus subgenus *Sarbecovirus*. Evolutionary Biology Available from: http://biorxiv.org/lookup/doi/10.1101/2020.07.07.190546

Wilm A, Aw PPK, Bertrand D, Yeo GHT, Ong SH, Wong CH, Khor CC, Petric R, Hibberd ML, Nagarajan N. 2012. LoFreq: a sequence-quality aware, ultra-sensitive variant caller for uncovering cell-population heterogeneity from high-throughput sequencing datasets. Nucleic Acids Research 40:11189–11201.

Worobey M, Pekar J, Larsen BB, Nelson MI, Hill V, Joy JB, Rambaut A, Suchard MA, Wertheim JO, Lemey P. 2020. The emergence of SARS-CoV-2 in Europe and North America. Science:eabc8169.

Wu A, Peng Y, Huang B, Ding X, Wang X, Niu P, Meng J, Zhu Z, Zhang Z, Wang J, et al.2020. Genome Composition and Divergence of the Novel Coronavirus (2019-nCoV) Originating in China. Cell Host & Microbe 27:325–328.

Wu F, Zhao S, Yu B, Chen Y-M, Wang W, Song Z-G, Hu Y, Tao Z-W, Tian J-H, Pei Y-Y, et al. 2020. A new coronavirus associated with human respiratory disease in China. Nature 579:265–269.

Yang Z. 1995. A Space-Time Process Model for the Evolution of DNA Sequences. Genetics 139:993–1005.

Yi Y, Lagniton PNP, Ye S, Li E, Xu R-H. 2020. COVID-19: what has been learned and to be learned about the novel coronavirus disease. Int. J. Biol. Sci. 16:1753–1766.

Yuen K-S, Ye Z-W, Fung S-Y, Chan C-P, Jin D-Y. 2020. SARS-CoV-2 and COVID-19: The most important research questions. Cell Biosci 10:40.

Yurkovetskiy L, Wang X, Pascal KE, Tomkins-Tinch C, Nyalile T, Wang Y, Baum A, Diehl WE, Dauphin A, Carbon\e C, et al. 2020. Structural and Functional Analysis of the D614G SARS-CoV-2 Spike Protein Variant. Cell, in press. Available from: https://www.cell.com/cell/fulltext/S0092-8674(20)31229-0

Zecha J, Lee C-Y, Bayer FP, Meng C, Grass V, Zerweck J, Schnatbaum K, Michler T, Pichlmair A, Ludwig C, et al. 2020. Data, Reagents, Assays and Merits of Proteomics for SARS-CoV-2 Research and Testing. Mol Cell Proteomics 19:1503–1522.

Zhou W, Chen T, Zhao H, Eterovic AK, Meric-Bernstam F, Mills GB, Chen K. 2014. Bias from removing read duplication in ultra-deep sequencing experiments. Bioinformatics 30:1073–1080.

Zhou P, Li H, Wang H, Wang L-F, Shi Z. 2012. Bat severe acute respiratory syndrome-like coronavirus ORF3b homologues display different interferon antagonist activities. Journal of General Virology 93:275–281.

